# Determining predictive metabolomic biomarkers of meniscal injury in dogs with cranial cruciate ligament disease

**DOI:** 10.1101/2022.08.22.504770

**Authors:** Christine R. Pye, Daniel C. Green, James R. Anderson, Marie M. Phelan, Eithne J. Comerford, Mandy J. Peffers

**Author notes:** Corresponding author, Corresponding author address: Christine Pye, Institute of Life Course and Medical Sciences, 1^st^ Floor William Henry Duncan Building, 6 West Derby Street, Liverpool, L7 8TX.

## Abstract

**Objectives:** The objective of this study was to use for the first time proton nuclear magnetic resonance spectroscopy (^1^H NMR) to examine the metabolomic profile of stifle joint synovial fluid from dogs with cranial cruciate ligament rupture with and without meniscal injuries. We hypothesised this would identify biomarkers of meniscal injury.

**Methods:** Stifle joint synovial fluid was collected from dogs undergoing stifle joint surgery or arthrocentesis for lameness investigations at three veterinary hospitals in the North-West of England. Samples underwent ^1^H NMR spectroscopy and metabolite identification. We used multivariate and univariate statistical analysis to identify differences in the metabolomic profile between dogs with cranial cruciate ligament rupture and meniscal injury, cranial cruciate ligament rupture without meniscal injury, and neither cranial cruciate ligament rupture nor meniscal injury, taking into consideration specific clinical variables.

**Results:** 154 samples of canine synovial fluid were included in the study. 64 metabolites were annotated to the ^1^H NMR spectra. Six spectral regions were found to be significantly altered between groups with cranial cruciate ligament rupture with and without meniscal injury, including three attributed to NMR mobile lipids (mobile lipid -CH_3_ [p=0.016], mobile lipid -n(CH_3_)_3_ [p=0.017], mobile unsaturated lipid [p=0.031]).

**Clinical Significance:** We identified an increase in NMR mobile lipids in the synovial fluid of dogs with meniscal injury which are of interest as potential biomarkers of meniscal injury, as well as understanding the metabolic processes that occur with meniscal injury.

## Introduction

Cranial cruciate ligament rupture (CCLR) is one of the most common causes of pelvic limb lameness in dogs (Witsberger *et al*., 2008). It presents a significant cause of morbidity amongst the canine population, and dogs with CCLR account for 0.56% of all cases presented to primary care veterinary practices in the UK (Taylor-Brown *et al*., 2015). One sequelae of joint instability caused by a loss of CCL function is tears to the menisci, found to occur in approximately 50% of cases at time of CCLR surgery (Bennett and May, 1991). The menisci are a pair of C shaped fibrocartilaginous structures located between the tibial plateau and femoral condyles (Kambic and McDevitt, 2005). They have several important functions including load bearing, load distribution and shock absorption, as well as contributing to joint stability, proprioception and joint lubrication (Arnoczky et al., 1980, Pozzi et al., 2010). Meniscal injuries as a result of CCLR most commonly affect the medial meniscus, due to its firmer attachment to the tibia caudally making it more prone to becoming trapped between the tibia and femur during cranial translocation of the tibia in stifle joints without a functioning CCL (Pozzi et al., 2008). Currently, treatment of meniscal injuries in dogs involves removal of part, or all, of the affected meniscus via an arthrotomy or arthroscopy (Franklin *et al*., 2017). The resultant loss of the normal meniscal structure leads to alterations in pressure distribution across the articular cartilage that can perpetuate the development of osteoarthritis (OA) (Pozzi *et al*., 2008).

Meniscal injuries can occur post-operatively after CCLR surgery, due to either residual joint instability, or failure to diagnose at the time of surgery (Metelman *et al*., 1995). The prevalence of these late meniscal injuries varies from 2.8% to 13.8% (Metelman et al., 1995, Fitzpatrick and Solano, 2010). Late meniscal injuries can be a cause of recurring stifle joint pain and lameness and are challenging for the veterinary practitioner to diagnose (Dillon *et al*., 2014). Affected dogs often present with a recurring lameness on the operated limb weeks or months after CCLR surgery, with clinical examination potentially revealing pain on stifle flexion, and/or a “click” on stifle flexion (Dillon *et al*., 2014, Case *et al*., 2008). The presence of a meniscal click has been found to be an unreliable diagnostic sign (McCready and Ness, 2016). Radiographs, useful in ruling out other causes of recurring lameness post-operatively, cannot show meniscal injuries directly. Won *et al*. (2020) recently investigated using radiographic joint space width as an indicator of meniscal injuries in lateral projections of the canine stifle, but this was only 40.5% sensitive. Further diagnostic imaging techniques for late meniscal injuries include low field or high field magnetic resonance imaging (MRI), computed tomography (CT) with arthrography, or ultrasound examination (McCready and Ness, 2016). Depending on the study, the sensitivity of these techniques in diagnosing meniscal injuries in dogs his 64-100% for low field MRI (Böttcher *et al*., 2010, Gonzalo-Orden *et al*., 2001) and 75-100% (Olive *et al*., 2014, Blond *et al*.,2008) for high field MRI. CT with arthrography and ultrasonography have sensitivities of 71% (Samii *et al*., 2009) and 90% (Mahn *et al*., 2005) respectively. All of these imaging techniques require either expensive specialised equipment, and/or advanced technical expertise, limiting the availability of these diagnostics in veterinary practice, and amount to a considerable cost. Surgical methods of diagnosis include either stifle joint arthroscopy (Van Gestel, 1985) or arthrotomy (Fitzpatrick and Solano, 2010). These surgical diagnostic hold inherent risk, although they also allow for treatment of any meniscal injuries at time of diagnosis (Ritzo *et al*., 2014). Furthermore, using surgery as a means of diagnosis has the risk of the animal undergoing an unnecessary surgical procedure if no meniscal injury is found (Blond *et al*., 2008).

Currently, there are no biomarkers of meniscal injury that can be used as a diagnostic aid. Also, no biomarkers of CCLR exist that could lead to earlier intervention or target preventative treatment in “at risk” stifles, such as the contralateral stifles of high-risk breeds. One potential source of biomarkers of stifle joint pathologies is synovial fluid (SF) (Boffa et al., 2020). SF is a viscous fluid, that is a dialysate of plasma (Ropes *et al*., 1940), and functions as a joint lubricant (Ghosh, 1994). SF has been found to contain a unique source of biomarkers of joint disease, due to its close proximity to structures within joints, (de Bakker et al., 2017, Anderson et al., 2018b). Previous studies have investigated potential cytokine and protein biomarkers of CCLR within canine stifle joint SF, including interleukin-8 (IL-8) (de Bruin *et al*., 2007b), anti-collagen type 1 antibodies (de Bruin *et al*., 2007a), matrix metalloproteinases (MMP) 2 and MMP9 (Boland *et al*., 2014, Murakami *et al*., 2016) and lubricin (Wang *et al*., 2020). There are relatively few studies examining biomarkers specific to meniscal injury within SF in any species. In humans, a recent study by Clair *et al*. (2019) found IL-6, monocyte chemoattractant protein-1 (MCP-1), macrophage inflammatory protein-1β (MIP-1β) and MMP-3 to be increased in the SF of humans with meniscal injuries compared to healthy controls.

Metabolomics allows the identification and quantification of small molecule metabolites and analysis of metabolic pathways within a variety of biofluids, cells and tissues (Bujak *et al*., 2015). Proton nuclear magnetic resonance (^1^H NMR) is a tool for metabolomics studies, having the benefits of being rapid, non-destructive and relatively inexpensive compared to other metabolomics tools such as mass spectrometry (Clarke *et al*., 2020). ^1^H NMR has been used successfully to investigate changes in the SF metabolomic profile in humans and horses with joint pathologies such as rheumatoid arthritis, OA, and septic arthritis (Anderson *et al*., 2018a, Anderson *et al*., 2018b, Clarke *et al*., 2020). Therefore, there is promise for using NMR spectroscopy to investigate biomarkers of joint pathology within canine SF. To date, only four published peer-reviewed studies have used NMR spectroscopy for metabolomic studies of canine stifle joint SF (Damyanovich *et al*., 1999a, Damyanovich *et al*., 1999b, Marshall *et al*., 2000, Crovace *et al*., 2006). However, these studies typically involved a small sample size, and were mainly focused on biomarkers of OA. NMR metabolomics could be used as a tool to study metabolites acting as biomarkers of meniscal injuries in CCLR affected stifle joints. This information could then potentially allow for the development of a simple, minimally invasive diagnostic test more reliable at detecting meniscal injuries, and late meniscal injuries, than pre-existing non-surgical diagnostic techniques.

We therefore hypothesise that the metabolomic profile of canine stifle joint SF will alter depending on the presence of CCLR, and depending on the presence of concurrent meniscal injuries.

## Materials and methods

### Ethical approval

Ethical approval for the collection of canine SF for use in this study was granted by the University of Liverpool Veterinary Research Ethics Committee (VREC634) as surplus clinical waste under the generic approval RETH00000553.

### Synovial fluid collection

Canine SF was collected from dogs undergoing surgery for CCLR or patella luxation, or as excess clinical waste from dogs undergoing arthrocentesis as part of lameness investigations from March 2018 to June 2021. Cases were recruited from three veterinary practices in the north-west of England, namely the University of Liverpool Small Animal Teaching Hospital, Leahurst, and the Animal Trust CIC Veterinary Practices at Bolton and Blackburn. Owners of the dogs were approached at time of admission for the procedure, and informed consent was obtained from the owners of the animals for the collection of the SF and participation in the project. SF was collected by stifle joint arthrocentesis as per the BSAVA guide to procedures in small animal practice (Bexfield and Lee, 2014). A 21 gauge to 23-gauge needle attached to a two to five millilitre sterile syringe (depending on the size of the dog) was inserted into the stifle joint space either medially or laterally to the patella ligament after sterile preparation of the skin, prior to first surgical incision. After aspiration of the SF, samples were placed in sterile 1.5ml Eppendorf tubes (Eppendorf UK Ltd, Stevenage, UK), and immediately refrigerated at 4°C.

### Synovial fluid processing

SF samples were transported on ice from the veterinary practices to the laboratory at the University of Liverpool, Leahurst Campus within 48 hours of collection. Samples stored for longer than 48 hours before processing were excluded from the study based on previous unpublished data examining metabolomic changes in the synovial fluid with elongated refrigerated storage time (Pye, 2021). Samples were centrifuged at 2540*g* at 4°C for 5 min. The supernatant was pipetted into 200 μl aliquots, and snap frozen in liquid nitrogen before storing at −80°C (Anderson *et al*., 2020).

### Clinical information on the canine participants

Clinical information regarding the dogs was collected. This included breed, age, sex and neuter status, body weight, body condition score (Laflamme, 1997), presence and degree of CCLR (whether partial or complete CCLR), presence of meniscal injury, location and type of meniscal injury (Bennett and May, 1991), presence of patella luxation, length of time of lameness, co-morbidities, medication being received by the dog and radiographic level of OA using two separate scoring systems (Innes *et al*.,2004, Wessely *et al*., 2017).

### NMR Metabolomics

#### Sample preparation for NMR metabolomics

SF samples were thawed on ice immediately prior to sample preparation for NMR spectroscopy. 100μL of each thawed SF sample was diluted to a final volume containing 50% (v/v) SF, 40% (v/v) dd 1H2O (18.2 MΩ), 100 mM PO43– pH 7.4 buffer (Na2HPO4, VWR International Ltd., Radnor, Pennsylvania, USA and NaH2PO4, Sigma-Aldrich, Gillingham, UK) in deuterium oxide (2H2O, Sigma-Aldrich) and 0.0025% (v/v) sodium azide (NaN3, Sigma-Aldrich). Samples were vortexed for 1 min, centrifuged at 13 000g and 4 °C for 5 min and 180μL transferred (taking care not to disturb any pelleted material) into 3 mm outer diameter NMR tubes using a glass pipette.

#### NMR metabolomics spectral acquisition

Spectra were acquired using a 700MHz Bruker Avance III spectrometer (Bruker Corporation, Billerica, Massachusetts, USA) with associated triple resonance inverse (TCI) cryoprobe and chilled Sample Jet auto-sampler. Software used for spectral acquisition and processing were Topspin 3.1 (Bruker Corporation, Billerica, Massachusetts, USA) and IconNMR 4.6.7 (Bruker Corporation).

1D ^1^H NMR spectra were acquired using a Carr-Purcell-Meiboom-Gill (CPMG) filter to suppress background signal disturbances from proteins and other endogenous constituents, and allow acquisition of small molecule metabolite signals (Carr and Purcell, 1954, Meiboom and Gill, 1958). A CPMG-1r vendor pulse sequence was used to achieve this. Water suppression was carried out by pre-saturation (Hoult, 1976). The CPMG spectra were acquired at 37 °C with a 15ppm spectral width, a four second interscan delay and 32 transients (Anderson *et al*., 2020).

#### NMR metabolomics spectral quality control

1D ^1^H NMR spectra were individually assessed to ensure minimum reporting standards were met (Sumner *et al*., 2007). Any samples that were deemed to have failed quality control were re-ran on the spectrometer up to a maximum of three spectral acquisitions. Any that failed after the third spectral acquisition were excluded from the study.

#### Metabolite annotation and identification

The NMR spectra were divided into spectral regions (termed “bins”) using Topspin 3.1 (Bruker Corporation, Massachusetts, USA), with each bin representing either single metabolite peaks or multiple metabolite peaks where peaks overlapped on the spectra. These bins were also examined using TameNMR (hosted by www.galaxy.liv.ac.uk). Bins were altered accordingly upon visualising the fit to the overlaid spectra to ensure the area under the peak was represented by the bin.

Metabolites were annotated to the spectra using Chenomx NMR Suite Profiler version 7.1 (Chenomx, Edmonton, Canada), a reference library of 302 mammalian metabolite NMR spectra. When metabolite peaks overlapped, multiple metabolites were annotated to the bin. When peaks were unable to be annotated to a metabolite, they were classed as being an “unknown” metabolite. Previous literature specifying metabolite chemical shifts and spectral appearance were examined to aid annotation of unknown areas. Downstream unique peak metabolite identification and in-house NMR metabolite standards were examined to confirm metabolite identities where possible.

A pattern file was created of the spectral bins and metabolites annotated to that bin. This is a spreadsheet outlining the bin boundaries in ppm, and the metabolites annotated to that bin (Pattern file included in supplementary material Table S1). The pattern file and the Bruker spectra files were input into TameNMR (galaxy.liv.ac.uk), in order to create a spreadsheet of binned spectra, with the relative intensities of each bin for each sample, which could then be used for statistical analysis of the spectra.

### Statistical analysis

#### Differences in clinical features of the canine participants

Analysis of the differences in clinical features between the groups in terms of age, sex and neuter status, body condition score, and radiographic OA score was undertaken using one-way analysis of variance (ANOVA) with Benjamini-Hochberg false discovery rate (FDR) adjustment, and significance set at p<0.05. Where any variable did not fit to ANOVA assumptions of having a common variance, Brown-Forsythe and Welch ANOVA tests with Benjamini-Hochberg FDR adjustment was carried out. These analyses and creation of graphs to visualise this data was carried out using GraphPad Prism 9.1.0 (GraphPad Software, San Diego, CA, USA).

#### Metabolomics data analysis

Metabolomics data was normalised using probabilistic quotient normalisation (PQN) (Dieterle *et al*., 2006), and Pareto scaled using R prior to statistical analysis (R Core Team, 2020). Unsupervised multivariate analysis was carried out using principal component analysis (PCA) on the normalised and scaled data using R. The variance between canine phenotypes was investigated through analysis of principal components 1 through 10 using One-Way ANOVAs or linear models depending on the data type. Briefly, CCLR, sex, neuter status, BCS, radiographic OA score and batch were numerically encoded and assessed against each principal component using a One-Way ANOVA. Age, Length of time of lameness, weight, length of time of storage pre-processing which were already numeric variables were assessed against each principal component using a linear model. All p values were corrected using FDR (Bejamini Hochberg) correction. Correlation matrices between phenotypes were computed using the spearmans correlation using the *cor* function in R and visualised using a heatmap generated with the *pheatmap* function in R (Kolde, 2012).

Univariate analysis was carried out using One-Way ANOVAs and One-Way analysis of co-variance (ANCOVAs) using R. To account for multiple testing across all 236 metabolite bins FDR correction was applied to the F-Test p value of each metabolite, significance was accepted at p < 0.05. For metabolites with an FDR < 0.05 Tukey’s honest significant difference post-hoc test was applied to assess between group variances. Age adjusted One-Way ANCOVAs were applied to each metabolite to assess differences between meniscal and no meniscal injury groups, FDR adjustment was applied as a above. Boxplots to visualise the changes in metabolite abundances were created in MetaboAnalyst 5.0 (https://www.metaboanalyst.ca), a software based on a metabolomics data analysis package written in R (the MetaboAnalystR package) (Pang *et al*., 2021).

## Results

### Clinical features of the canine participants

For the metabolomic study, 191 samples of canine stifle joint SF were collected and submitted for NMR spectroscopy. Of these, 14 samples had been stored for longer than 48 hours prior to collection for processing, and were subsequently excluded from the study. Four samples were from cases in which the meniscal injury status was unknown, and were also excluded from the study. Nineteen samples were excluded as they failed to meet minimum reporting standards (Sumner *et al*., 2007) after three spectral acquisitions.

In total, 154 canine stifle joint SF samples were included in the statistical analysis. These were divided into three groups, namely CCLR without meniscal injury (n=72), CCLR with meniscal injury (n=65), and control group with neither CCLR or meniscal injury present (n=17). The control group consisted of 13 cases of patella luxation, three cases from arthrocentesis of the stifle joints during lameness investigations which subsequently were found to have no pathology, and one sample from a case with fraying of the caudal cruciate ligament. Information regarding the signalment of the dogs in each group is shown in Table 1.

**Table 1.**
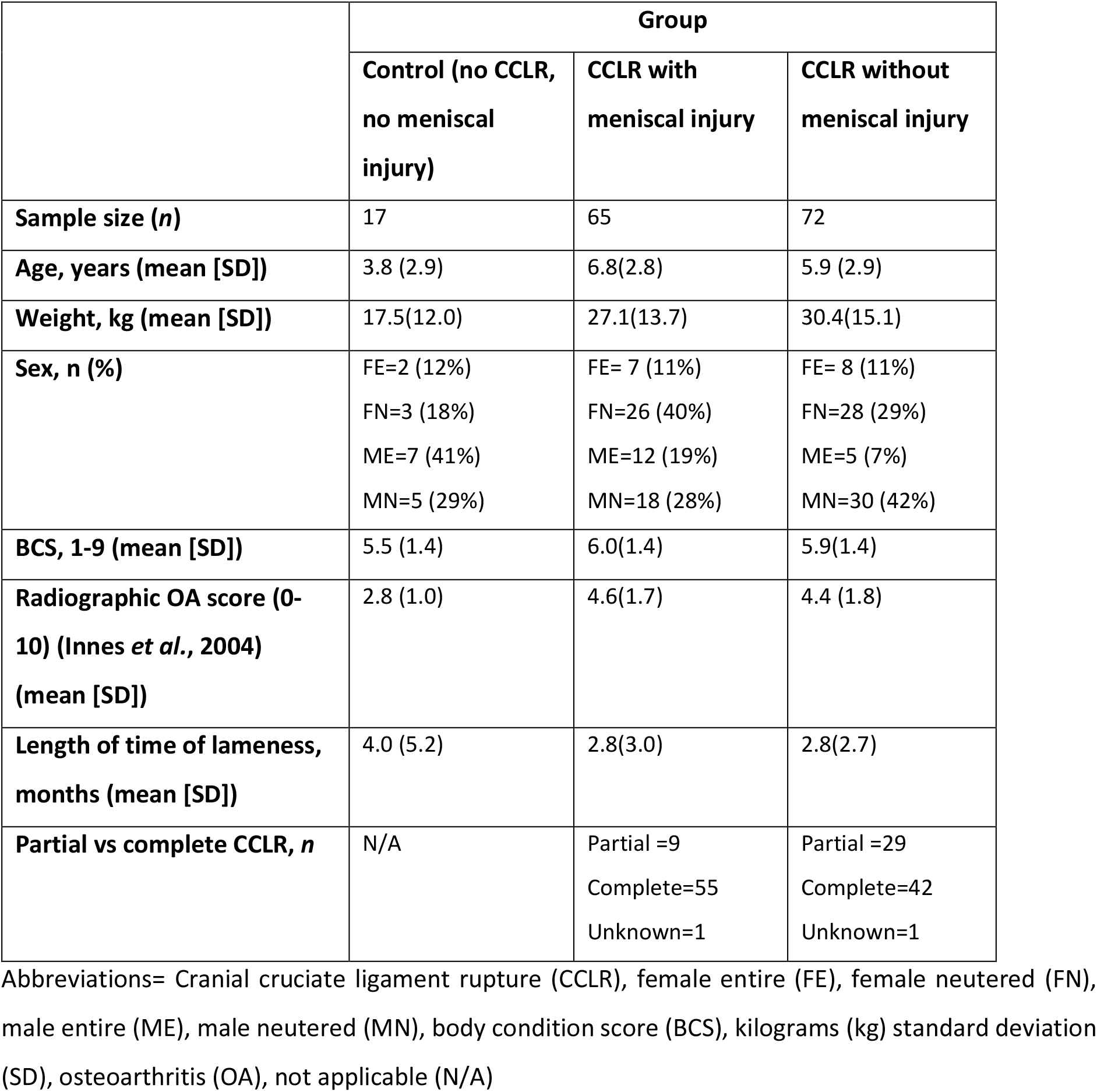
Clinical features of the canine participants included in the nuclear magnetic resonance metabolomic study of biomarkers of meniscal injury in canine stifle joint synovial fluid with cranial cruciate ligament rupture.

|  | Group |  |  |
| --- | --- | --- | --- |
|  | Control (no CCLR, no meniscal injury) | CCLR with meniscal injury | CCLR without meniscal injury |
| **Sample size (n)** | 17 | 65 | 72 |
| **Age, years (mean [SD])** | 3.8 (2.9) | 6.8(2.8) | 5.9 (2.9) |
| **Weight, kg (mean [SD])** | 17.5(12.0) | 27.1(13.7) | 30.4(15.1) |
| **Sex, n (%)** | FE=2 (12%)<br>FN=3 (18%)<br>ME=7 (41%)<br>MN=5 (29%) | FE= 7 (11%)<br>FN=26 (40%)<br>ME=12 (19%)<br>MN=18 (28%) | FE= 8 (11%)<br>FN=28 (29%)<br>ME=5 (7%)<br>MN=30 (42%) |
| **BCS, 1-9 (mean [SD])** | 5.5 (1.4) | 6.0(1.4) | 5.9(1.4) |
| **Radiographic OA score (0-10) (Innes *et al.*, 2004) (mean [SD])** | 2.8 (1.0) | 4.6(1.7) | 4.4 (1.8) |
| **Length of time of lameness, months (mean [SD])** | 4.0 (5.2) | 2.8(3.0) | 2.8(2.7) |
| **Partial vs complete CCLR, n** | N/A | Partial =9<br>Complete=55<br>Unknown=1 | Partial =29<br>Complete=42<br>Unknown=1 |

Abbreviations= Cranial cruciate ligament rupture (CCLR), female entire (FE), female neutered (FN), male entire (ME), male neutered (MN), body condition score (BCS), kilograms (kg) standard deviation (SD), osteoarthritis (OA), not applicable (N/A)

There was a significant difference between the control group and the CCLR groups with or without meniscal injury in terms of age, weight, and radiographic OA score, but not with BCS (Figure 1). There was no significant difference between groups CCLR with meniscal injury and CCLR without meniscal injury in terms of these clinical variables, although age was closest to reaching significance between the two groups (p=0.07, Mean Difference=0.86 [0.01 to 1.73 95% CI]).

**Figure 1.**
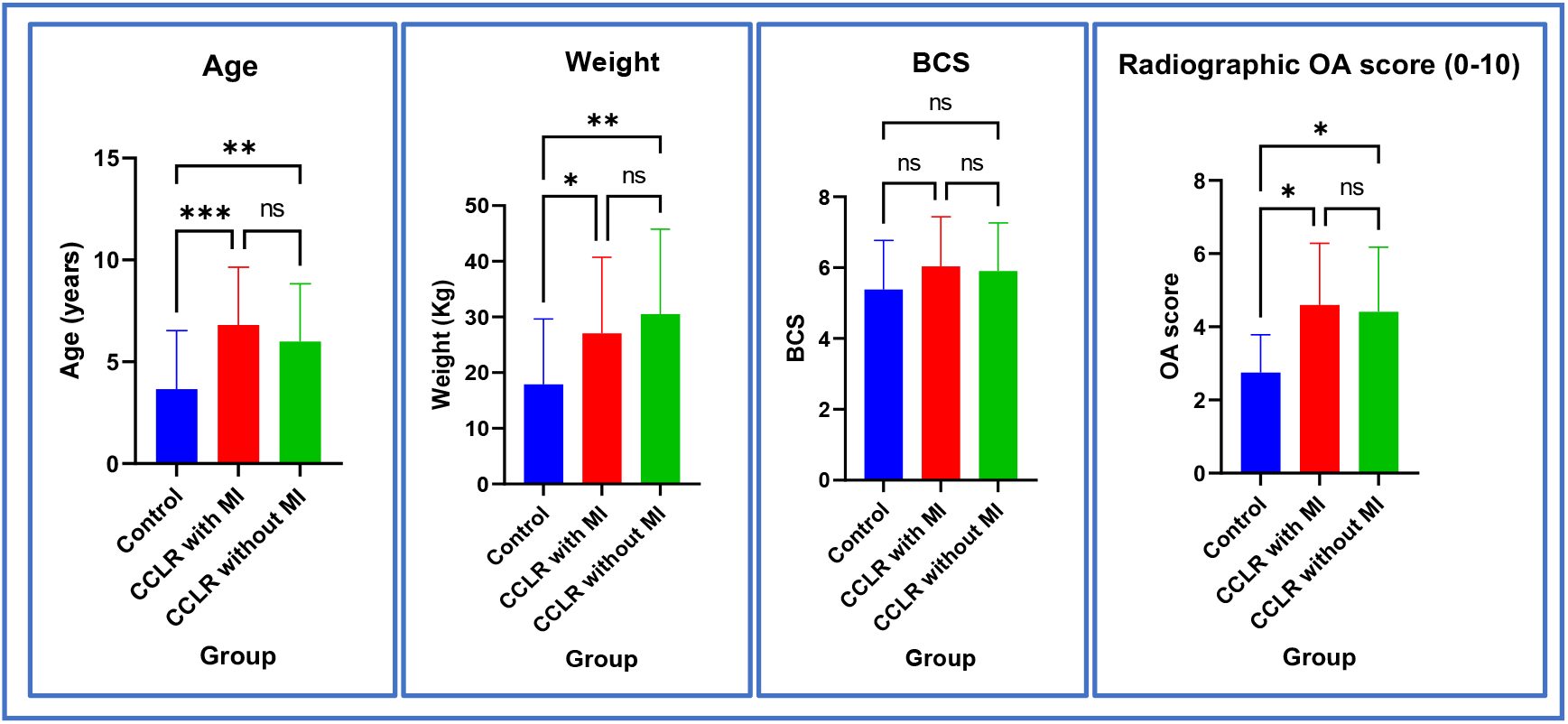
Clinical features of the canine participants between groups. Column Bar graphs represent mean and standard deviation. Groups= Control (n=17), CCLR with meniscal injury (n=65), CCLR without meniscal injury (n=72). Significance testing performed with either one-way ANOVA, or Brown-Forsythe and Welch ANOVA (depending on whether data had common variance) with Benjamini-Hochberg false discovery rate adjustment (CCLR=cranial cruciate ligament disease, MI=meniscal injury, OA=osteoarthritis, ns=not significant, *=p<0.05, **=p<0.01, ***=p<0.001).

### Metabolite annotation and identification

Spectra were divided into 246 bins. Of these, 84 (34%) remained with an unknown metabolite identification, and 162 (66% of bins) were annotated to one or more metabolites. In total, 64 metabolites were annotated to the spectra (Table 2). The pattern file of these bins and metabolite annotations is included in the supplementary information (Table S1). Any bins containing ethanol peaks were excluded from the statistical analysis, due to ethanol being considered a contaminant in NMR, usually either during the collection of the SF from the sterilisation of skin with surgical spirit, or during the processing steps (van der Sar *et al*., 2015). Propylene glycol, a metabolite found in solvents used in pharmaceuticals (Zar *et al*., 2007) was found in one spectrum, and so those bins were excluded so as to not bias the statistical analysis.

**Table 2.** Metabolites annotated to canine stifle joint synovial fluid nuclear magnetic resonance spectra, including HMDB identification number where possible, and level of identification according to the Metabolomics Standard Initiative (Sumner *et al*., 2007).

| AMINO ACIDS |  |  | FATTY AND ORGANIC ACIDS |  |  |
| --- | --- | --- | --- | --- | --- |
| Metabolite name | HMDB number | MSI ID Level | Metabolite name | HMDB number | MSI ID Level |
| 2-AMINOADIPATE | HMDB0000510 | Level 2 | 2-HYDROXYVALERATE | HMDB0001863 | Level 2 |
| BETAINE | HMDB0000043 | Level 2 | 2-METHYLGLUTARATE | HMDB0000422 | Level 2 |
| CREATINE | HMDB0000064 | Level 1 | 2-PHENYLPROPIONATE | HMDB0011743 | Level 2 |
| CREATINE PHOSPHATE | HMDB0001511 | Level 2 | 3- HYDROXYISOVALERATE | HMDB0000754 | Level 2 |
| CREATININE | HMDB0000562 | Level 1 | 4-PYRIDOXATE | HMDB0000017 | Level 2 |
| CREATININE PHOSPHATE | HMDB0041624 | Level 2 | AZELATE | HMDB0000784 | Level 2 |
| L-ALANINE | HMDB0000161 | Level 1 | CITRATE | HMDB0000094 | Level 1 |
| L-ALLOISOLEUCINE | HMDB0000557 | Level 2 | FORMATE | HMDB0000142 | Level 2 |
| L-GLUTAMINE | HMDB0000641 | Level 1 | GLYCOCHOLATE | HMDB0000138 | Level 2 |
| L-HISTIDINE | HMDB0000177 | Level 1 | GLYCOLATE | HMDB0000115 | Level 2 |
| L-ISOLEUCINE | HMDB0000172 | Level 1 | GLYCYLPROLINE | HMDB0000721 | Level 2 |
| L-LEUCINE | HMDB0000687 | Level 1 | ISOBUTYRIC ACID | HMDB0001873 | Level 2 |
| L-LYSINE | HMDB0000182 | Level 1 | L-CARNITINE | HMDB0000062 | Level 2 |
| L-METHIONINE | HMDB0000696 | Level 1 | L-GLUTAMATE | HMDB0060475 | Level 2 |
| L-PHENYLALANINE | HMDB0000159 | Level 1 | LACTATE | HMDB0000190 | Level 1 |
| L-THREONINE | HMDB0000167 | Level 1 | METHYLSUCCINATE | HMDB0001844 | Level 2 |
|  |  |  | MOBILE LIPIDS | N/A | Level 3 |
| L-TYROSINE | HMDB0000158 | Level 1 | PYRUVATE | HMDB0000243 | Level 1 |
| L-VALINE | HMDB0000883 | Level 1 | SEBACATE | HMDB0000792 | Level 2 |
|  |  |  | THYMOL | HMDB0001878 | Level 2 |
| SUGARS |  |  | OTHERS |  |  |
| Metabolite name | HMDB number | MSI ID Level | Metabolite name | HMDB number | MSI ID Level |
| D-GALACTOSE | HMDB0000143 | Level 2 | 3-HYDROXY-3-METHYLGLUTARATE | HMDB0041199 | Level 2 |
| D-GLUCOSE | HMDB0000122 | Level 1 | 3-METHYLHISTIDINE | HMDB0000479 | Level 2 |
| FRUCTOSE | HMDB0000660 | Level 2 | ACETAMINOPHEN | HMDB0001859 | Level 2 |
| GLUCITOL | HMDB0000247 | Level 2 | ACETONE | HMDB0001659 | Level 2 |
| MANNITOL | HMDB0000765 | Level 2 | ACETYLCHOLINE | HMDB0000895 | Level 2 |
| MANNOSE | HMDB0000169 | Level 1 | CHOLINE | HMDB0000097 | Level 1 |
| MYOINOSITOL | HMDB0000211 | Level 1 | DIMETHYL SULFONE | HMDB0004983 | Level 2 |
|  |  |  | DTTP | HMDB0001342 | Level 2 |
|  |  |  | ETHANOL | HMDB0000108 | Level 1 |
|  |  |  | HISTAMINE | HMDB0000870 | Level 2 |
|  |  |  | O-CRESOL | HMDB0002055 | Level 2 |
|  |  |  | P-CRESOL | HMDB0001858 | Level 2 |
|  |  |  | PROPYLENE GLYCOL | HMDB0001881 | Level 2 |
|  |  |  | SN-GLYCERO-3-PHOSPHOCHOLINE | HMDB0000086 | Level 2 |
|  |  |  | TAU-METHYLHISTIDINE | HMDB0000479 | Level 2 |
|  |  |  | TRIGONELLINE | HMDB0000875 | Level 2 |
|  |  |  | XANTHINE | HMDB0000292 | Level 2 |

Abbreviations: HMDB=Human metabolome database; MSI= Metabolomics standards initiative

### Metabolomic statistical analysis results

#### Multivariate analysis of canine synovial fluid metabolome with respect to CCLR and meniscal injury status

Multivariate PCA was undertaken to compare the differences in the overall metabolome between the groups (no CCLR and no meniscal injury [control group], CCLR with meniscal injury, CCLR without meniscal injury) (Figure 2). Over PC one and two, there were clustering of the groups, indicating little overall difference in the metabolome over these PCs (Figure 2a). Associations between different phenotypes of the canine participants and PC one to ten found that PC three and four were primarily associated with CCLR and meniscal injury (supplementary material Figure S1). PCA of the groups plotted over PC three and four showed some separation of the control group with the groups CCLR with and without meniscal injury, indicating that the control group appears to have a differing metabolome from the other two groups (Figure 2b). The differences in the metabolome based on the presence of either a partial or complete tear found no significant differences in the metabolome.

**Figure 2.**
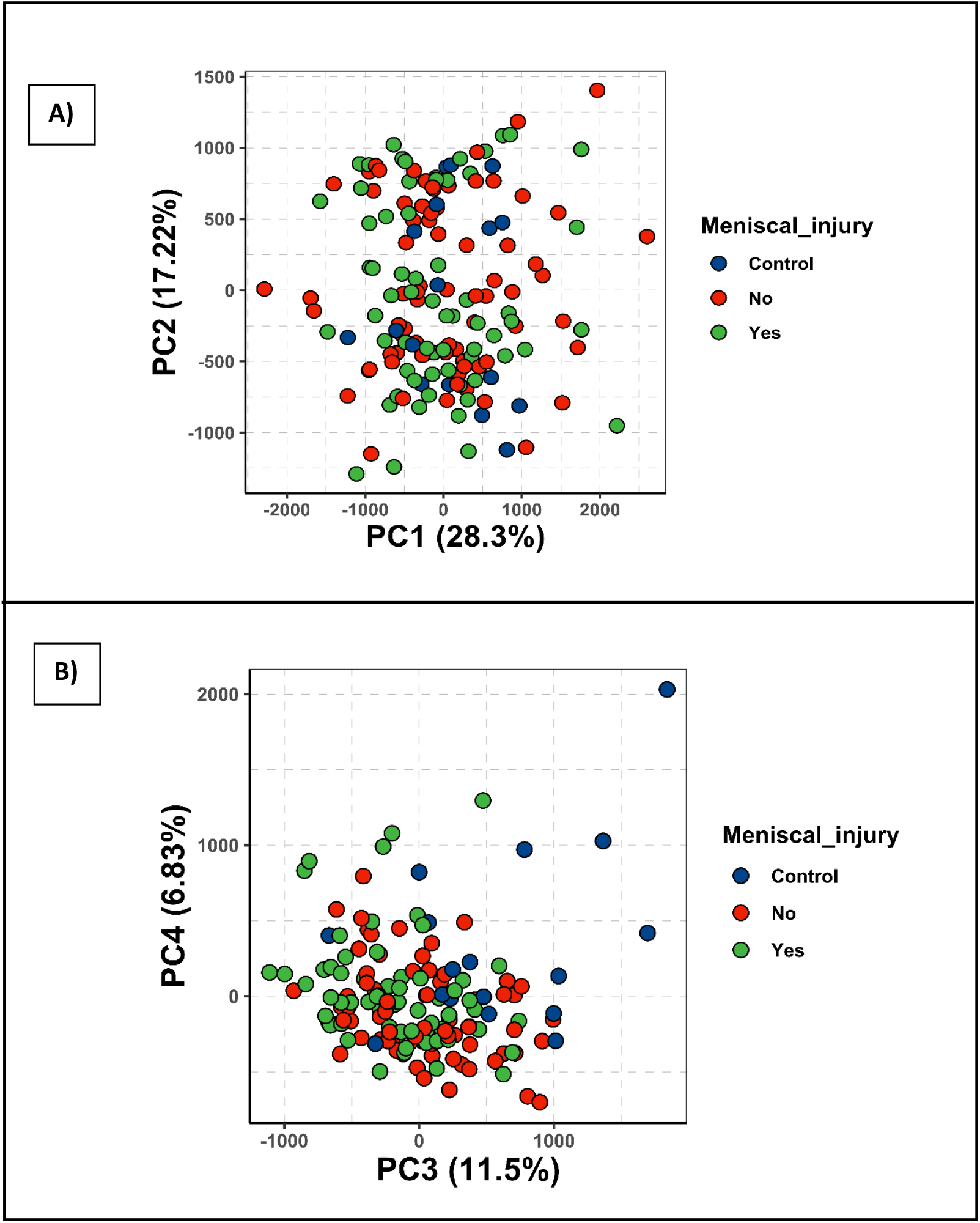
Principal component analysis 2D scores plot of samples of canine stifle joint synovial fluid following NMR. Meniscal injury status plotted over A) PC1 and PC2 and B) PC3 and PC4. Blue (control)=no CCLR, no meniscal injury; red (no)=CCLR without meniscal injury; green (yes)=CCLR with meniscal injury.

#### Univariate analysis of canine synovial fluid metabolome with respect to clinical variables, CCLR and meniscal injury status

Analysis of metabolite changes with respect to clinical variables also found significantly altered metabolites with differing weight (supplementary information Figure S2), age (supplementary information Figure S3) and radiographic OA score of the dogs (supplementary information Figure S4). It was therefore not possible to accurately assess the changes with regard to CCLR (irrespective of the meniscal injury status) versus the control group, due to the significant differences between these clinical variables in the control group versus the other groups, and the small sample size in the control group compared to the other groups.

Univariate analysis of metabolomic differences between the three groups (no CCLR and no meniscal injury [control group], CCLR with meniscal injury and CCLR without meniscal injury) was then undertaken to examine metabolomic changes in the presence of meniscal injury within the SF. There were six spectral bins that were below the threshold of significance (p<0.05), and two others that neared the threshold (p<0.06) after one-way ANOVA testing with FDR adjusted p-values and Tukey’s HSD *post-hoc* test (Table 3).

**Table 3.** Metabolites found to be significantly altered in canine stifle joint synovial fluid between those dogs with CCLR with meniscal injury and those with CCLR without meniscal injury using ANOVA testing.

| Bin number | Chemical shift (ppm) | Metabolite(s) annotated to bin | Mean difference (RI) | 95% CI | FDR adjusted p-value |
| --- | --- | --- | --- | --- | --- |
| 145 | 3.268-3.272 | UNKNOWN | -46.57 | -80.45 to -12.69 | 0.004 |
| 230 | 1.071-1.080 | METHYLSUCCINATE and/or 2-METHYLGLUTARATE | 21.97 | 5.91 to 38.04 | 0.004 |
| 129 | 3.362-3.371 | METHANOL | -40.04 | -74.27 to -5.80 | 0.017 |
| 210 | 1.936-2.020 | GLYCYLPROLINE, ISOLEUCINE and UNKNOWN | 37.96 | 2.79 to 73.12 | 0.031 |
| 152 | 3.203-3.238 | MOBILE LIPID - N(CH<sub>3</sub>)<sub>3</sub> | 104.42 | 4.85 to 203.98 | 0.037 |
| 246 | 0.789-0.891 | MOBILE LIPID -CH<sub>3</sub> | 82.25 | 3.37 to 161.13 | 0.039 |
| 37 | 5.212-5.353 | MOBILE UNSATURATED LIPID | 42.04 | -0.06 to 84.14 | 0.050 |
| 224 | 1.199-1.312 | MOBILE LIPID -(CH<sub>2</sub>)<sub>n</sub> | 88.78 | -2.63 to 180.19 | 0.059 |

Abbreviations: ppm= parts per million; RI=relative intensity, CI= confidence interval; FDR= false discovery rate

As it was noted that mobile lipids were also significantly altered with increasing age of the canine participants (supplementary information Figure S3), and that groups CCLR with meniscal injury and CCLR without meniscal injury had a slight, although insignificant (p=0.07, Mean Difference=0.86 years [0.01 to 1.73 95% CI]) difference in terms of age of the canine participants in each group, ANCOVAs were undertaken to control for age within the samples. The results of these ANCOVAs controlling for age are shown in Table 4. After controlling for age, three out of four spectral regions annotated to mobile lipids had an increased significant difference between groups CCLR with meniscal injury and CCLR without meniscal injury (Figure 3). Another mobile lipid signal lost significance, as did the spectral bin annotated to methylsuccinate and/or 2-methylglutarate. A complete list of the ANCOVA outputs are included in the supplementary information (Table S2).

**Table 4.** Metabolites found to be significantly altered in canine stifle joint synovial fluid between those dogs with CCLR with meniscal injury and those with CCLR without meniscal injury using ANCOVA testing after controlling for age of the dogs.

| Bin number | Chemical shift (ppm) | Metabolite(s) annotated to bin | Mean difference | 95% CI | FDR adj p-value |
| --- | --- | --- | --- | --- | --- |
| 145 | 3.268-3.272 | UNKNOWN | 46.94 | 18.6 to 75.3 | 0.004 |
| 129 | 3.362-3.371 | METHANOL | 40.01 | 11.3 to 68.7 | 0.009 |
| 246 | 0.789-0.891 | MOBILE LIPID -CH<sub>3</sub> | -78.88 | -142.84 to -14.91 | 0.016 |
| 152 | 3.203-3.238 | MOBILE LIPID -N(CH<sub>3</sub>)<sub>3</sub> | -99.38 | -179.03 to -19.73 | 0.017 |
| 210 | 1.936-2.020 | GLYCYLPROLINE, ISOLEUCINE and UNKNOWN | -36.35 | -64.7 to -7.97 | 0.019 |
| 37 | 5.212-5.353 | MOBILE UNSATURATED LIPID | -40.06 | -73.96 to -6.16 | 0.031 |

Abbreviations: ppm= parts per million; CI= confidence interval; FDR= false discovery rate

**Figure 3.**
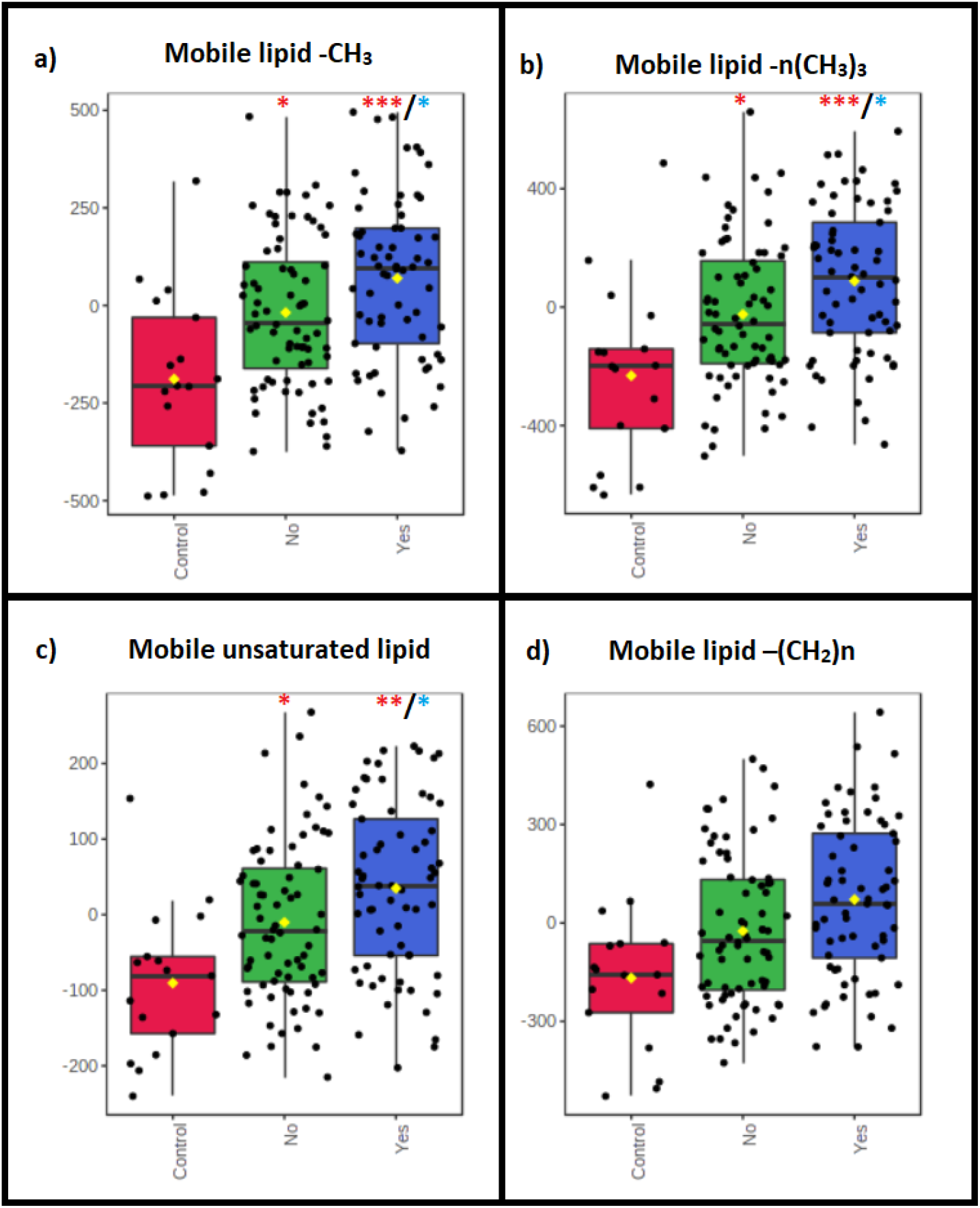
Altered mobile lipids on ^1^H NMR with respect to meniscal injury status in canine stifle joint synovial fluid from dogs. Bar plots on the left of each panel show the original abundances (mean +/− SD), and box and whisker plots on the right show the normalised abundances. Control (red)= no CCLR or meniscal injury. No (green)= CCLR without meniscal injury. Yes (blue)= CCLR with meniscal injuy. Red stars above boxplots denote significance against control group. Blue stars above boxplots denote significance against ‘No’ group. Significance testing was completed using one-way ANCOVAs controlling for age of the canine participants in each group (*=p<0.05, **=p<0.01, ***=p<0.0001).

## Discussion

A consistent finding when analysing the metabolomic differences in groups based on meniscal injury status was an increase in mobile lipids on the ^1^H NMR spectra in stifle joints with meniscal injury. Mobile lipids are NMR lipid resonances that arise from isotropically tumbling, relatively non-restricted molecules such as methyl and methylene resonances belonging to lipid acyl chains (Hakumäki and Kauppinen, 2000, Delikatny *et al*., 2011). These arise primarily from triglycerides, fatty acids and cholesteryl esters in lipid droplets, and also from phospholipidic acyl chains if not embedded in lipid membrane bilayers (Mannechez *et al*., 2005). Lipids serve various important functions in biological systems, including as components of cell membranes and other cellular organelles, acting as an energy source, and having a crucial role in signalling and regulation of cellular processes (Onal *et al*., 2017). Many biological processes have been associated with changes in NMR mobile lipids, including cell necrosis and apoptosis, malignancy, inflammation, proliferation and growth arrest (Hakumäki and Kauppinen, 2000). Lipid analysis of SF in humans have found differential abundance of lipids with different disease states, including OA, rheumatoid arthritis and trauma (Wise *et al*., 1987). A more recent NMR lipidomic study in SF from canine and human OA affected joints found an increase in numerous lipid species in OA compared to healthy controls in both species (Kosinska *et al*., 2016).

There are a number of possible hypotheses for the increase in mobile lipid resonances found in the SF of dogs with CCLR and concurrent meniscal injury compared to CCLR without meniscal injury in this study. Injury to the meniscus could lead to ‘damage to cellular phospholipid membranes, resulting in the release of lipids into the SF. Human menisci have also been found to contain lipid debris that could have an impact on SF lipid concentrations in meniscal injury (Ghadially and Lalonde, 1981). Also, lipid droplets could be released from the intracellular environment due to cell necrosis or apoptosis in the damaged meniscal tissue (Uysal *et al*., 2008), leading to an increased concentration of lipid droplets in the SF. Lipid droplets have been found to play a key role in inflammation, as such it may be that meniscal tears lead to a release of lipid droplets to facilitate in the inflammatory response within the joint (Melo *et al*., 2011). As lipid droplets contain mediators of inflammation such as pro-inflammatory cytokines, lipids could also potentiate inflammatory changes in meniscal injury affected joints (Ichinose *et al*., 1998). Alterations in SF lipid composition and lipid species can also have a role in affecting the lubricating ability of the SF (Antonacci *et al*., 2012). The concentration of phospholipid species in human SF have been found to be increased in OA affected joints, therefore the observed increase in lipids could also be an attempt to improve lubrication of the SF after meniscal injury in order to have protective effects on the articular cartilage (Kosinska *et al*., 2015).

Amongst the other differentially abundant metabolites between groups, CCLR with and without meniscal injury, was methanol. Although methanol could be considered a contaminant in NMR (Fulmer *et al*., 2010), it has also been found to be a naturally occurring metabolite in humans, either through dietary consumption in various fruit and vegetables, the artificial sweetener aspartame, alcohol, or through actions of gut microbiota (Dorokhov *et al*., 2015). Some of these sources cannot be ruled out, and therefore the decision not to remove methanol from analysis was made. However, its association with meniscal injury remains unclear. Unfortunately, the spectral bin that had the highest significance in differential abundance between CCLR with and without meniscal injury SF groups was unable to be identified using reference libraries and in-depth literature searches.

One of the spectral bins that also showed a significant increase in canine SF in dogs with CCLR and meniscal injury compared to CCLR without meniscal injury was a region that had overlapping NMR peaks annotated to glycylproline, isoleucine, and an unknown metabolite. This region also requires further work to confirm the identity of the specific metabolites attributed to this area. It is possible that in this region at 1.93 to 2.02 ppm there were also mobile lipid resonances, as fatty acyl chains have been noted to be attributed to this region (Delikatny *et al*., 2011). This would correlate with the findings of increases in mobile lipids with meniscal injury. Further work to identify the metabolites in this region could include undertaking a 2D NMR experiment, or a “spiking” experiment involving adding known concentrations of specific metabolites to the sample (Dona *et al*., 2016)

There were a number of metabolite peaks that are, as yet, unidentified on the canine SF spectra, including one that was found to be significantly altered with meniscal injury. Further work is required in identifying these regions, such as undertaking a 2D NMR experiment, or spiking experiment, as mentioned above. Alternatively, using more sensitive methods of molecule identification, such as mass spectrometry, could improve the number of metabolite identifications in the samples.

One of the limitations of our study was the lack of a balanced control group to compare with the CCLR affected joints. There are several reasons for this. Firstly, collection of “normal” SF via arthrocentesis from joints without pre-existing pathology involves a level of risk, including introducing infection into the joint, and the need for sedation or anaesthetic for the protocol (Bexfield and Lee, 2014). Therefore, this would have ethical implications, and was outside the ethical approval for this study. SF from dogs with no stifle joint pathology collected post-mortem would have been subjected to metabolite changes that would have compromised the comparison to the diseased groups (Donaldson and Lamont, 2015). Control samples in this study were collected from dogs undergoing surgery for patella luxation, or excess SF from dogs undergoing arthrocentesis from investigations of lameness. These were cases without CCLR or meniscal injuries, but also may not have been completely without pathological changes, as patella luxation can be cause of OA and synovitis (Roush, 1993). Patella luxation also tends to be more common in smaller breeds of dogs, and as primarily a congenital disease, cases often cases show clinical signs of lameness at a younger age than CCLR affected dogs (LaFond *et al*., 2002, Rudd Garces *et al*., 2021). Both these factors lent towards the control group being on average younger and smaller than the CCLR groups, with less osteoarthritic changes. This, along with the fewer samples collected in the time constraints of this study affected the ability to infer conclusions from the metabolite changes between the control and other groups in terms of CCLR alone. The inclusion of a more donors in the control group of healthy, non-diseased canine stifle joint SF would be of value in future work to allow analysis of changes due to CCLR and OA in the canine stifle joint.

There were factors such as diet and level of exercise that have been found to affect the metabolome of human serum that were not been accounted for in this study (Esko *et al*., 2017, Sakaguchi *et al*., 2019). However, unlike humans, dogs tend to have a less variable diet, and also exercise is likely to be similar between the canine participants, as the standard advice for dogs affected by CCLR is to limit exercise. Medications were found to be too heterogeneous between the dogs in this study from which to make any statistical conclusions but are known to affect the metabolomic profile of biofluids (Um *et al*., 2009).

This study is the first of its kind in using ^1^H NMR spectroscopy to identify biomarkers of meniscal injury with SF. SF lipid species appear to be of interest in the study of biomarkers of meniscal injury, and future work to identify the lipid species involved by undertaking a lipidomics experiment, such as NMR using lipid extracts from the SF samples, or using light chromatography coupled with mass spectrometry (LC-MS) lipidomics. This further work could prove useful in exploring the potential for a diagnostic marker of meniscal injury in canine SF.

The work presented here is also of translational value into metabolomics studies in human and other mammalian species. No SF biomarker has been found to date in human SF with meniscal injury, therefore this research could also lead to the investigation of biomarkers of meniscal injury in human SF.

## Acknowledgements

We would like to acknowledge and give thanks to all the staff at the University of Liverpool Small Animal Teaching Hospital and the Animal Trust for their help in collecting samples for this study, and to all the owners who gave their consent for their dogs to be included in the study. Our thanks also go to veterinary students Matt Fitzgerald and Callum Burke for their preliminary studies, and to Alex Simon for his work on the radiographic OA scoring. Finally, we thank BSAVA PetSavers for providing funding that allowed this study to happen.

