## Supplementary information for "Determining predictive metabolomic biomarkers of meniscal injury in dogs with cranial cruciate ligament disease"

Figures

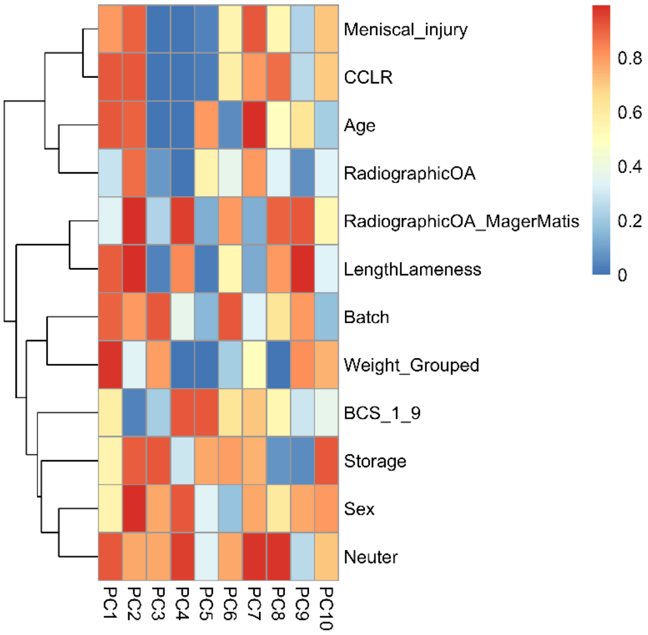

**Figure S1. Heat map showing association of clinical features of dogs whose synovial fluid was submitted for metabolomic analysis in the study with the first ten principal components in a principal component analysis.** The key on the right denotes FDR corrected p-values from 0 (blue) to 1 (red). Meniscal injury and cranial cruciate ligament rupture (CCLR) appear to be associated primarily with principal components (PC) three to five. (OA= osteoarthritis, BCS= body condition score, Radiographic OA= global assessment of radiographic OA score (0-3), RadiographicOA_MagerMatis= Mager and Matiss *et al.* radiographic osteoarthritis score (0-45), LengthLameness= length of time of lameness, Storage= time stored at four degrees Celsius prior to processing and freezing).

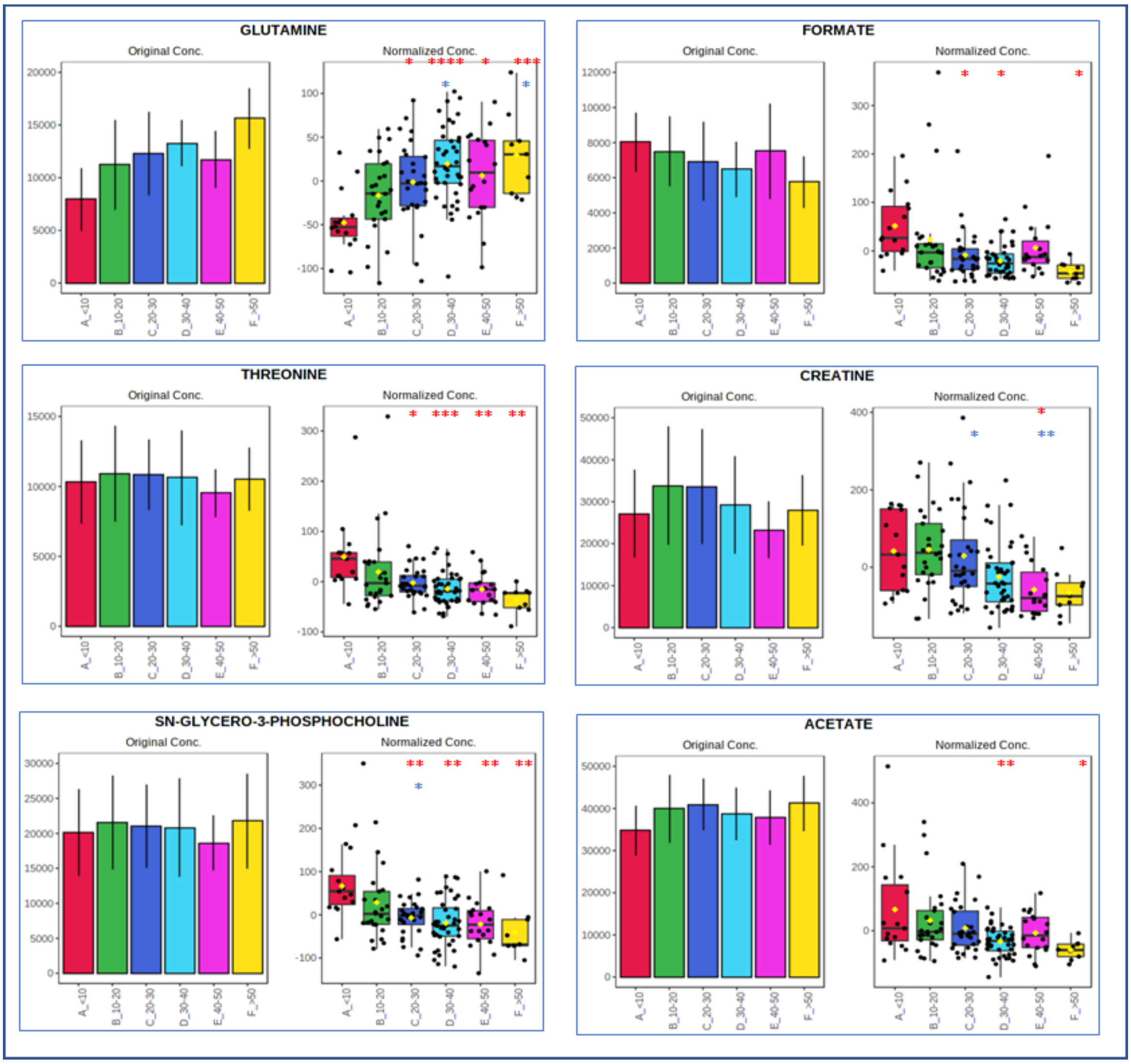

**Figure S2. Bar plots and box and whisker plots showing changes in metabolites with increasing body weight in canine stifle joint synovial fluid from dogs with cranial cruciate ligament rupture.** Bar plots on the left show the original values (mean +/- SD), and box and whisker plots on the right show the normalised values. The x axis shows the weight groups in Kg. Key to colours of bar charts: Red= <10kg, Green = 10-20kg, Navy blue= 20-30kg, Light blue= 30-40kg, Pink=40-50kg, Yellow= >50kg. Red stars above the boxplots denote significance in comparison with the <10kg group, blue stars above the box plots denote significance in comparison with the 10-20kg group; *=p<0.05, **=p<0.01, ***=p<0.001, ****=p<0.0001.

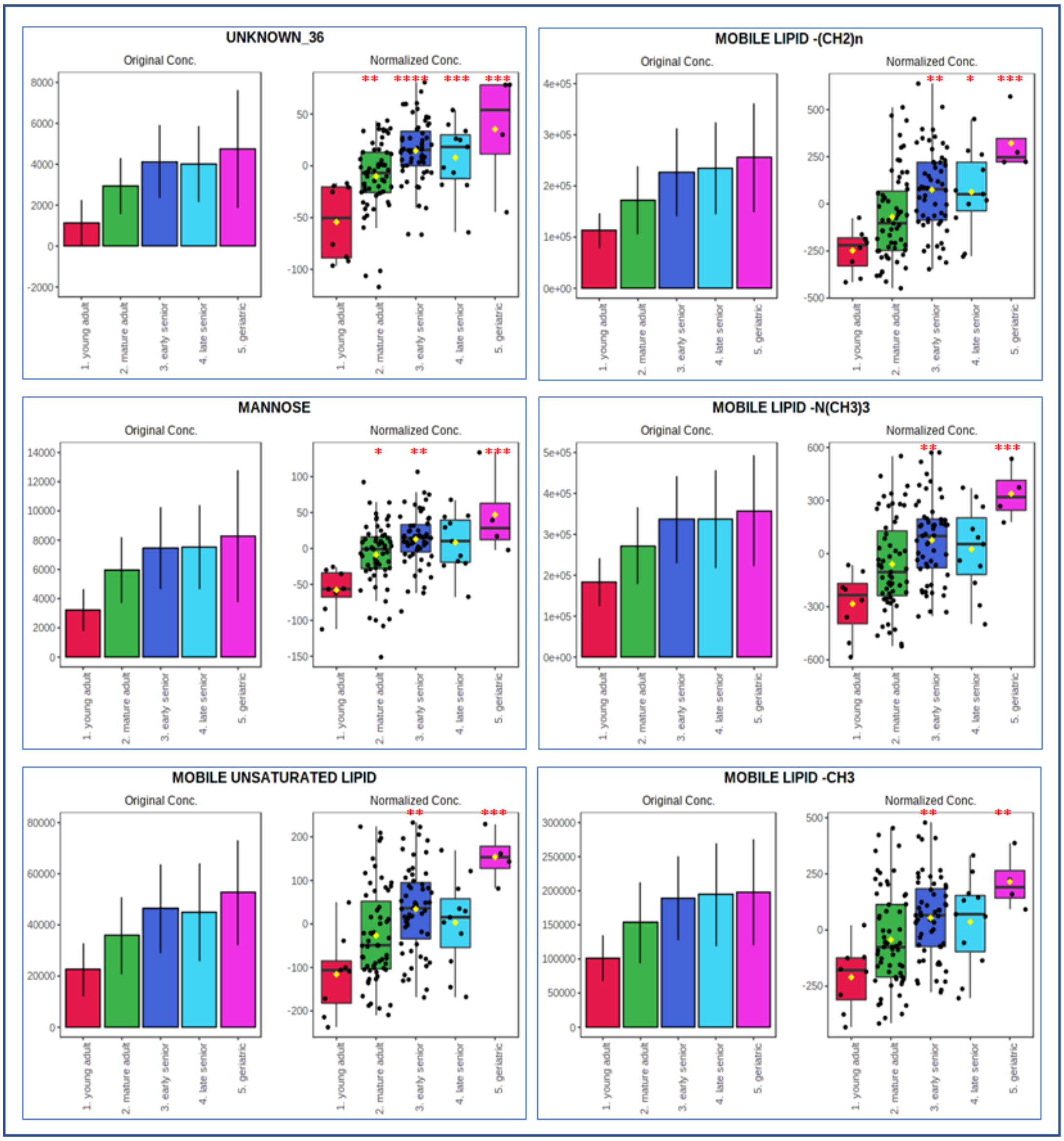

**Figure S3. Bar plots and box and whisker plots showing changes in metabolites with increasing age of the dog in canine stifle joint synovial fluid from dogs with cranial cruciate ligament rupture.** Bar plots on the left show the original values (mean +/- SD), and box and whisker plots on the right show the normalised values. The x axis shows the age groups from young adult to geriatric. Key to colours of bar charts: Red=young adult, Green =mature adult, Navy blue=early senior, Light blue=late senior, Pink=geriatric. Red stars above boxplots denote significance in comparison with group 1 (young adult); *=p<0.05, **=p<0.01, ***=p<0.001, ****=p<0.0001.

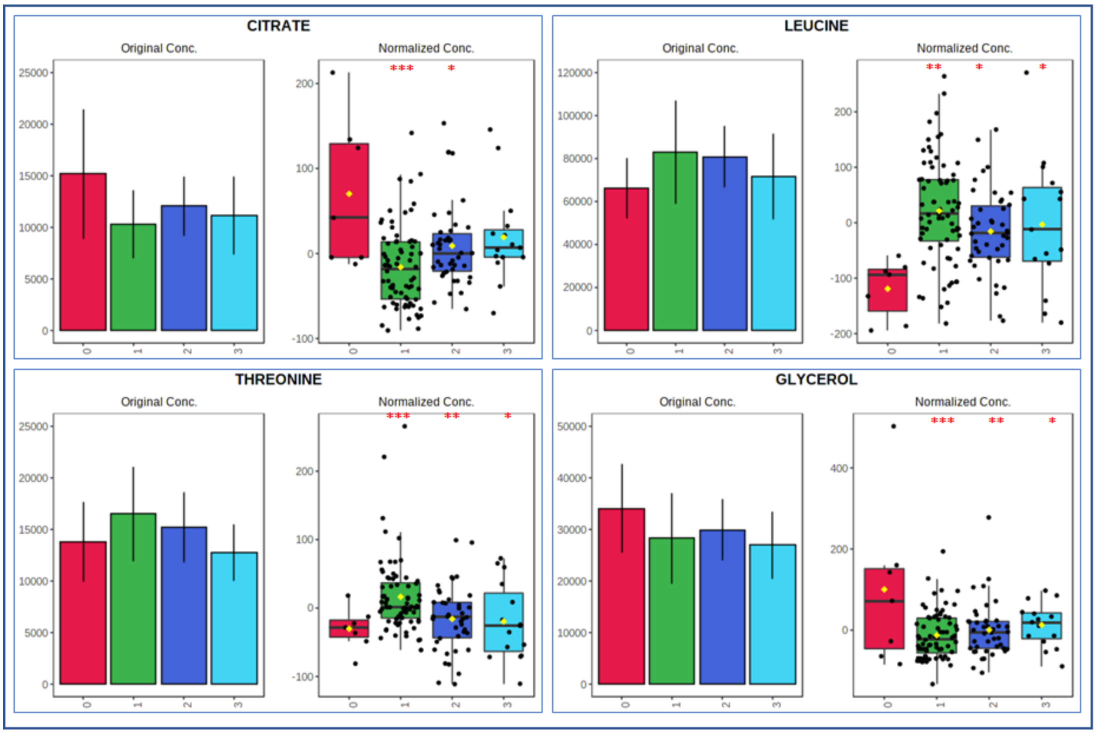

**Figure S4. Bar plots and box and whisker plots showing changes in metabolites with increasing radiographic osteoarthritis score of the dog in canine stifle joint synovial fluid from dogs with cranial cruciate ligament rupture.** Bar plots on the left show the original values (mean +/- SD), and box and whisker plots on the right show the normalised values. The x axis shows OA scores from 0-3 (Red=0, Green=1, Navy blue=2, Light blue=3). Stars above boxplots denote significance against OA score 0 (*=p<0.05, **=p<0.01, ***=p<0.001).

Tables

Table S1. Pattern file for canine synovial fluid $^{1}H$ 1D NMR spectra, showing the boundaries of each spectral bin in parts per million and the metabolite(s) annotated to each bin.

| Higher boundary (ppm) | Lower boundary (ppm) | Metabolite(s) Annotated and bin number |
|---|---|---|
| 8.4669 | 8.4456 | FORMATE_1 |
| 7.9301 | 7.9154 | XANTHINE_2 |
| 7.9153 | 7.8321 | 4-PYRIDOXATE_ANSERINE_3 |
| 7.8309 | 7.7756 | HISTIDINE_4 |
| 7.7325 | 7.7083 | dTTP_t-methylhistidine_5 |
| 7.6489 | 7.613 | UNKNOWN_6 |
| 7.6129 | 7.5826 | UNKNOWN_7 |
| 7.5609 | 7.5265 | UNKNOWN_8 |
| 7.4536 | 7.4376 | PHENYLALANINE_9 |
| 7.4375 | 7.4259 | PHENYLALANINE_10 |
| 7.4258 | 7.4149 | PHENYLALANINE_11 |
| 7.4148 | 7.3979 | UNKNOWN_12 |
| 7.3978 | 7.3823 | PHENYLALANINE_2-PHENYLPROPRIONATE_13 |
| 7.3822 | 7.3711 | PHENYLALANINE_2-PHENYLPROPRIONATE_14 |
| 7.371 | 7.3559 | PHENYLALANINE_2-PHENYLPROPRIONATE_15 |
| 7.3475 | 7.3336 | PHENYLALANINE_16 |
| 7.3335 | 7.3205 | PHENYLALANINE_17 |
| 7.3204 | 7.3115 | UNKNOWN_18 |
| 7.2786 | 7.2336 | UNKNOWN_19 |
| 7.2166 | 7.1978 | TYROSINE_20 |
| 7.1977 | 7.1823 | TYROSINE_21 |
| 7.1674 | 7.153 | UNKNOWN_22 |
| 7.1529 | 7.138 | UNKNOWN_23 |
| 7.1324 | 7.0929 | UNKNOWN_24 |
| 7.0837 | 7.0525 | HISTIDINE_t-METHYLHISTIDINE_25 |
| 7.0348 | 6.9856 | UNKNOWN_26 |
| 6.9201 | 6.9056 | TYROSINE_ACETAMINOPHEN_O-CRESOL_GLUTAMINE_27 |
| 6.9055 | 6.889 | TYROSINE_ACETAMINOPHEN_O-CRESOL_GLUTAMINE_28 |
| 6.8493 | 6.7967 | P-CREOSOL_29 |
| 6.6654 | 6.6331 | UNKNOWN_30 |
| 6.5255 | 6.5171 | UNKNOWN_31 |
| 6.2689 | 6.2621 | DTTP_32 |
| 5.919 | 5.9002 | UNKNOWN_33 |
| 5.8253 | 5.8022 | UNKNOWN_34 |
| 5.4086 | 5.3953 | UNKNOWN_35 |
| 5.3663 | 5.3532 | UNKNOWN_36 |
| 5.3531 | 5.251 | MOBILE UNSATURATED LIPID_37 |
| 5.2509 | 5.2115 | GLUCOSE_38 |

| | | |
|---|---|---|
| 5.1968 | 5.1783 | MANNOSE_39 |
| 5.124 | 5.1094 | UNKNOWN_40 |
| 5.1093 | 5.0981 | UNKNOWN_41 |
| 5.098 | 5.0883 | UNKNOWN_42 |
| 4.9087 | 4.9019 | MANNOSE_43 |
| 4.4626 | 4.454 | TRIGONELLINE_44 |
| 4.3912 | 4.3821 | UNKNOWN_45 |
| 4.382 | 4.3702 | UNKNOWN_46 |
| 4.3353 | 4.281 | SN-GLYCERO-3-PHOSPHOCHOLINE_47 |
| 4.2734 | 4.2649 | THREONINE_48 |
| 4.2648 | 4.2372 | THREONINE_49 |
| 4.2371 | 4.2258 | UNKNOWN_50 |
| 4.2257 | 4.2172 | UNKNOWN_51 |
| 4.2076 | 4.1831 | UNKNOWN_52 |
| 4.1487 | 4.1386 | UNKNOWN_53 |
| 4.1385 | 4.1327 | UNKNOWN_54 |
| 4.1326 | 4.0933 | LACTATE_55 |
| 4.0932 | 4.0871 | UNKNOWN_56 |
| 4.084 | 4.0778 | GALACTOSE_CHOLINE_57 |
| 4.0777 | 4.0733 | GALACTOSE_CHOLINE_58 |
| 4.0732 | 4.0667 | GALACTOSE_CHOLINE_59 |
| 4.0666 | 4.0603 | GALACTOSE_CHOLINE_60 |
| 4.0602 | 4.0489 | CREATININE_61 |
| 4.0208 | 4.0092 | UNKNOWN_62 |
| 4.0091 | 4.0046 | UNKNOWN_63 |
| 4.0045 | 3.999 | UNKNOWN_64 |
| 3.9989 | 3.9949 | UNKNOWN_65 |
| 3.9948 | 3.9869 | UNKNOWN_66 |
| 3.9868 | 3.983 | UNKNOWN_67 |
| 3.9829 | 3.9776 | UNKNOWN_68 |
| 3.9775 | 3.9664 | UNKNOWN_69 |
| 3.9663 | 3.9561 | UNKNOWN_70 |
| 3.956 | 3.9482 | GLYCOLATE_71 |
| 3.9481 | 3.9327 | UNKNOWN_72 |
| 3.9326 | 3.9258 | CREATINE_73 |
| 3.9257 | 3.9204 | UNKNOWN_74 |
| 3.9203 | 3.9052 | GLUCOSE_75 |
| 3.9026 | 3.8858 | GLUCOSE_BETAINE_76 |
| 3.8857 | 3.8724 | MANNITOL_77 |
| 3.8699 | 3.8639 | MANNITOL_78 |
| 3.8638 | 3.8604 | MANNITOL_79 |
| 3.8603 | 3.852 | GLUCOSE_80 |
| 3.8519 | 3.8466 | UNKNOWN_81 |
| 3.8465 | 3.8355 | GLUCOSE_82 |
| 3.8354 | 3.8321 | GLUCOSE_83 |
| 3.832 | 3.8283 | GLUCOSE_84 |

| | | |
|---|---|---|
| 3.8282 | 3.8251 | GLUCOSE_85 |
| 3.825 | 3.8212 | GLUCOSE_86 |
| 3.8211 | 3.8185 | GLUCOSE_87 |
| 3.8184 | 3.812 | GLUCOSE_MANNITOL_88 |
| 3.8119 | 3.8001 | ALANINE_MANNITOL_89 |
| 3.8 | 3.7907 | ALANINE_GLUTAMINE_90 |
| 3.7906 | 3.7808 | GLUCOSE_ALANINE_MANNITOL_GALACTOSE_GLUCITOL_91 |
| 3.7807 | 3.7752 | GLUCOSE_ANSERINE_GALACTOSE_LYCINE_MANNITOL_GLUCITOL_92 |
| 3.7751 | 3.7723 | ALANINE_GLUCOSE_GLUCITOL_MANNITOL_93 |
| 3.7722 | 3.7634 | GLUCOSE_GALACTOSE_LYCINE_GLUTAMATE_MANNITOL_94 |
| 3.7633 | 3.7552 | GLUCOSE_GALACTOSE_LYCINE_GLUTAMATE_GLUCITOL_MANNITOL_95 |
| 3.7502 | 3.7419 | GLUCOE_ALLOISOLEUCINE_2-AMINOADIPATE_GALACTOSE_O-ACETYLCHOLINE_LEUCINE_96 |
| 3.7418 | 3.7314 | GLUCOSE_O-ACETYLCHOLINE_2-AMINOADIPATE_GALACTOSE_LEUCINE_97 |
| 3.7313 | 3.7257 | GLUCOSE_FRUCTOSE_98 |
| 3.7256 | 3.7214 | GLUCOSE_99 |
| 3.7213 | 3.718 | GLUCOSE_PI-METHYLHISTIDINE_100 |
| 3.7179 | 3.7123 | UNKNOWN_101 |
| 3.7122 | 3.6989 | GLUCOSE_T-METHYLHISTIDINE_GALACTOSE_MANNITOL_102 |
| 3.6988 | 3.6917 | MANNITOL_103 |
| 3.6916 | 3.683 | MANNITOL_104 |
| 3.6829 | 3.6766 | MANNITOL_105 |
| 3.6765 | 3.6676 | ETHANOL_106 |
| 3.6675 | 3.662 | GLYCEROL_ETHANOL_107 |
| 3.6619 | 3.6582 | UNKNOWN_ETHANOL_108 |
| 3.6581 | 3.6521 | ETHANOL_109 |
| 3.652 | 3.6471 | GLYCEROL_ETHANOL_110 |
| 3.647 | 3.6437 | ETHANOL_111 |
| 3.6436 | 3.6368 | UNKNOWN_112 |
| 3.6367 | 3.6299 | MYO-INOSITOL_113 |
| 3.6298 | 3.6218 | UNKNOWN_114 |
| 3.6217 | 3.6176 | MYO-INOSITOL_VALINE_115 |
| 3.6175 | 3.6051 | VALINE_THREONINE_116 |
| 3.605 | 3.5921 | UNKNOWN_117 |
| 3.592 | 3.5691 | UNKNOWN_GLYCEROL_118 |
| 3.569 | 3.5598 | GLYCINE_119 |
| 3.5597 | 3.5558 | GLYCEROL_120 |
| 3.5557 | 3.5505 | GLUCOSE_121 |
| 3.5504 | 3.5433 | GLUCOSE_122 |
| 3.5432 | 3.5367 | GLUCOSE_123 |
| 3.5366 | 3.5281 | GLUCOSE_124 |
| 3.5185 | 3.5068 | GLUCOSE_125 |
| 3.5067 | 3.4487 | GLUCOSE_126 |
| 3.4486 | 3.4404 | UNKNOWN_127 |
| 3.4403 | 3.3815 | GLUCOSE_128 |
| 3.3706 | 3.3621 | METHANOL_129 |

| | | |
|---|---|---|
| 3.362 | 3.3578 | UNKNOWN_130 |
| 3.3577 | 3.3451 | UNKNOWN_131 |
| 3.345 | 3.3399 | UNKNOWN_132 |
| 3.3398 | 3.3324 | UNKNOWN_133 |
| 3.3323 | 3.3108 | UNKNOWN_134 |
| 3.3107 | 3.3057 | PI-METHYLHISTIDINE_135 |
| 3.3056 | 3.301 | UNKNOWN_136 |
| 3.2988 | 3.2919 | UNKNOWN_137 |
| 3.2918 | 3.28817 | UNKNOWN_138 |
| 3.2835 | 3.2741 | UNKNOWN_139 |
| 3.2832 | 3.276 | UNKNOWN_141 |
| 3.2759 | 3.2736 | UNKNOWN_142 |
| 3.274 | 3.2833 | UNKNOWN_140 |
| 3.2735 | 3.2738 | UNKNOWN_143 |
| 3.2727 | 3.2719 | UNKNOWN_144 |
| 3.2718 | 3.2679 | UNKNOWN_145 |
| 3.2678 | 3.2645 | GLUCOSE_BETAINE_146 |
| 3.2644 | 3.2608 | BETAINE_147 |
| 3.2607 | 3.2574 | UNKNOWN_BETAINE_GLUCOSE_148 |
| 3.2573 | 3.2491 | GLUCOSE_149 |
| 3.249 | 3.2443 | UNKNOWN_150 |
| 3.2442 | 3.2379 | GLUCOSE_151 |
| 3.2378 | 3.2033 | MOBILE LIPID -N(CH3)3_152 |
| 3.2032 | 3.1952 | CHOLINE_153 |
| 3.1951 | 3.1809 | UNKNOWN_154 |
| 3.1763 | 3.1274 | DIMETHYL SULFONE_155 |
| 3.1273 | 3.0975 | UNKNOWN_156 |
| 3.0518 | 3.0429 | CREATININE_CREATINE_LYSINE_TYROSINE_CREATININE-PHOSPHATE_157 |
| 3.0428 | 3.0369 | LYSINE_CREATININE_CREATINE-PHOSPHATE_CREATINE_TYROSINE_158 |
| 3.0368 | 3.0268 | LYSINE_159 |
| 3.0267 | 3.0173 | LYSINE_160 |
| 3.0172 | 3.0079 | UNKNOWN_161 |
| 3.0078 | 2.9992 | UNKNOWN_162 |
| 2.9991 | 2.989 | UNKNOWN_163 |
| 2.9678 | 2.9478 | UNKNOWN_164 |
| 2.9477 | 2.9061 | UNKNOWN_165 |
| 2.906 | 2.8951 | UNKNOWN_166 |
| 2.895 | 2.8849 | UNKNOWN_167 |
| 2.872 | 2.8316 | UNKNOWN_168 |
| 2.6866 | 2.6672 | UNKNOWN_169 |
| 2.6671 | 2.6522 | UNKNOWN_170 |
| 2.6521 | 2.6436 | UNKNOWN_171 |
| 2.6435 | 2.6304 | UNKNOWN_172 |
| 2.5733 | 2.5565 | UNKNOWN_173 |
| 2.5564 | 2.5357 | CITRATE_174 |
| 2.5356 | 2.5166 | CITRATE_175 |

| | | |
|---|---|---|
| 2.5165 | 2.5122 | UNKNOWN_176 |
| 2.5121 | 2.5063 | UNKNOWN_177 |
| 2.5062 | 2.5003 | UNKNOWN_178 |
| 2.5002 | 2.4803 | GLUTAMINE_179 |
| 2.4802 | 2.4693 | GLUTAMINE_180 |
| 2.4692 | 2.4571 | GLUTAMINE_181 |
| 2.457 | 2.4451 | GLUTAMINE_182 |
| 2.445 | 2.4344 | GLUTAMINE_183 |
| 2.4343 | 2.4229 | GLUTAMINE_184 |
| 2.4228 | 2.4085 | GLUTAMINE_185 |
| 2.4084 | 2.3964 | 3-HYDROXY-3-METHYLGLUTARATE_186 |
| 2.3963 | 2.3791 | DTTP_GLUTAMATE_ISOBUTYRATE_UNKNOWN_187 |
| 2.379 | 2.3714 | PYRUVATE_188 |
| 2.3713 | 2.3681 | GLUTAMATE_3-HYDROXYISOVALERATE_189 |
| 2.368 | 2.3352 | GLUTAMATE_190 |
| 2.2994 | 2.2869 | VALINE_191 |
| 2.2868 | 2.28 | ACETOACETATE_VALINE_SUCCINYLACETONE_192 |
| 2.2799 | 2.2706 | VALINE_UNKNOWN_193 |
| 2.2705 | 2.2609 | VALINE_2-AMINOADIPATE_P-CREOSOL_2-METHYLGLUTARATE_GLYCYLPROLINE_UNKNOWN_194 |
| 2.2608 | 2.2509 | 2-AMINOADIPATE_THYMOL_VALINE_2-METHYLGLUTARATE_GLYCYLPROLINE_VALINE_P-CRESOL_195 |
| 2.2508 | 2.2364 | UNKNOWN_196 |
| 2.2363 | 2.2294 | ACETONE_O-CRESOL_197 |
| 2.2062 | 2.1854 | UNKNOWN_198 |
| 2.1853 | 2.1726 | GLUTAMINE_2-METHYLGLUTARATE_AZELATE_SEBACATE_199 |
| 2.1725 | 2.168 | GLUTAMINE_ACETAMINOPHEN_200 |
| 2.1679 | 2.1614 | GLUTAMINE_ACETAMINOPHEN_METHIONINE_2-METHYLGLUTARATE_GLUTAMATE_201 |
| 2.1613 | 2.1517 | GLUTAMINE_GLUTAMATE_202 |
| 2.1516 | 2.1404 | METHIONINE_GLUTAMINE_GLUTAMATE_203 |
| 2.1403 | 2.0939 | GLUTAMINE_GLUTAMATE_204 |
| 2.0938 | 2.0708 | N-ACETYLCYSTEINE_GLUTAMATE_ALLOISOLEUCINE_205 |
| 2.0707 | 2.0638 | GLUTAMATE_206 |
| 2.0637 | 2.0542 | GLUTAMATE_207 |
| 2.0541 | 2.0315 | N-ACETYLGLUTAMINE_GLUTAMATE_UNKNOWN_208 |
| 2.0314 | 2.0202 | UNKNOWN_GLUTAMATE_209 |
| 2.0201 | 1.9363 | GLYCYLPROLINE_ISOLEUCINE_UNKNOWN_210 |
| 1.9272 | 1.9149 | ACETATE_211 |
| 1.9148 | 1.8691 | LYSINE_UKNOWN_212 |
| 1.8065 | 1.7592 | UNKNOWN_213 |
| 1.7591 | 1.7036 | LYCINE_LEUCINE_214 |
| 1.7035 | 1.6345 | LEUCINE_LYSINE_2-HYDROXYVALERATE_215 |
| 1.6344 | 1.5936 | 2-HYDROXYVALERATE_UNKNOWN_216 |
| 1.5935 | 1.5419 | UNKNOWN_217 |
| 1.5418 | 1.4975 | LYSINE_218 |
| 1.5141 | 1.4362 | ALANINE_219 |

| | | |
|---|---|---|
| 1.4296 | 1.4077 | 2-PHENYLPROPRIONATE_ALLOISOLEUCINE_220 |
| 1.4076 | 1.3774 | 2-HYDROXYVALERATE_ALLOISOLEUCINE_221 |
| 1.3773 | 1.3474 | UNKNOWN_222 |
| 1.3473 | 1.3167 | LACTATE_THREONINE_223 |
| 1.3166 | 1.1986 | MOBILE LIPID -(CH2)n (VLDL)_224 |
| 1.1985 | 1.1924 | ETHANOL_225 |
| 1.1889 | 1.1819 | ETHANOL_226 |
| 1.1818 | 1.1671 | UNKNOWN_227 |
| 1.153 | 1.1453 | PROPYLENE GLYCOL_228 |
| 1.1452 | 1.1361 | PROPYLENE GLYCOL_229 |
| 1.08 | 1.0717 | METHYLSUCCINATE_2-METHYLGLUTARATE_230 |
| 1.0716 | 1.0625 | METHYLSUCCINATE_2-METHYLGLUTARATE_231 |
| 1.0563 | 1.026 | VALINE_ISOBUTYRATE_232 |
| 1.0202 | 1.0087 | ISOLEUCINE_233 |
| 1.0086 | 1.0004 | ISOLEUCINE_234 |
| 1.0003 | 0.9943 | VALINE_235 |
| 0.9942 | 0.9905 | ALLOISOLEUCINE_VALINE_236 |
| 0.9904 | 0.9841 | VALINE_237 |
| 0.984 | 0.9778 | ALLOISOLEUCINE_VALINE_238 |
| 0.9777 | 0.9671 | LEUCINE_ALLOISOLEUCINE_239 |
| 0.967 | 0.9564 | LEUCINE_240 |
| 0.9563 | 0.9432 | LEUCINE_ISOLEUCINE_ALLOISOLEUCINE_241 |
| 0.9431 | 0.9334 | ALLOISOLEUCINE_ISOLEUCINE_242 |
| 0.9333 | 0.9219 | ISOLEUCINE_243 |
| 0.9175 | 0.9063 | GLYCOCHOLATE_244 |
| 0.9062 | 0.8977 | UNKNOWN_245 |
| 0.8908 | 0.7892 | MOBILE LIPID -CH3(VLDL)_246 |

Table S4. All outcomes of Tukey HSD post-hoc test from metabolites found to be significant between groups on ANCOVA.

| Group comparison | Bin number | Metabolite(s) annotated to bin | Difference in means | SE | 95% CI(low) | 95% CI(high) | FDR adjusted p-value | significance level |
|---|---|---|---|---|---|---|---|---|
| Control vs no | 180 | GLUTAMINE | -99.89 | 19.62 | -138.66 | -61.11 | 3.17E-06 | **** |
| | 181 | GLUTAMINE | -148.54 | 29.61 | -207.04 | -90.04 | 3.60E-06 | **** |
| | 179 | GLUTAMINE | -57.16 | 11.41 | -79.69 | -34.62 | 4.52E-06 | **** |
| | 182 | GLUTAMINE | -150.17 | 30.62 | -210.67 | -89.67 | 4.68E-06 | **** |
| | 120 | GLYCEROL | 131.66 | 27.94 | 76.47 | 186.86 | 8.28E-06 | **** |
| | 183 | GLUTAMINE | -97.10 | 20.56 | -137.72 | -56.48 | 9.63E-06 | **** |
| | 31 | UNKNOWN | 24.45 | 5.23 | 14.12 | 34.77 | 1.92E-05 | **** |
| | 184 | GLUTAMINE | -57.83 | 12.85 | -83.23 | -32.43 | 2.19E-05 | **** |
| | 227 | UNKNOWN | 171.21 | 37.07 | 97.97 | 244.45 | 2.47E-05 | **** |
| | 147 | BETAINE | 366.02 | 82.59 | 202.83 | 529.21 | 2.69E-05 | **** |
| | 200 | GLUTAMINE_ACETAMINOPHEN | -67.43 | 14.78 | -96.63 | -38.23 | 3.12E-05 | **** |
| | 189 | GLUTAMATE_3-HYDROXYISOVALERATE | 118.69 | 27.17 | 65.01 | 172.37 | 3.48E-05 | **** |
| | 190 | GLUTAMATE | 113.55 | 27.26 | 59.69 | 167.42 | 7.84E-05 | **** |
| | 223 | LACTATE_THREONINE | 324.03 | 75.67 | 174.51 | 473.55 | 9.87E-05 | **** |
| | 199 | GLUTAMINE_2-METHYLGLUTARATE_AZELATE_SEBACATE | -35.15 | 8.47 | -51.89 | -18.42 | 1.66E-04 | *** |
| | 204 | GLUTAMINE_GLUTAMATE | -74.23 | 18.14 | -110.07 | -38.39 | 2.08E-04 | *** |
| | 55 | LACTATE | 137.94 | 34.04 | 70.67 | 205.21 | 2.44E-04 | *** |
| | 3 | 4-PYRIDOXATE_ANSERINE | 17.57 | 4.55 | 8.57 | 26.56 | 2.52E-04 | *** |
| | 209 | UNKNOWN_GLUTAMATE | 93.52 | 23.59 | 46.90 | 140.14 | 3.41E-04 | *** |
| | 185 | GLUTAMINE | -37.40 | 9.99 | -57.14 | -17.65 | 3.89E-04 | *** |
| | 211 | ACETATE | 97.85 | 26.74 | 45.01 | 150.69 | 5.25E-04 | *** |
| | 134 | UNKNOWN | -28.55 | 7.91 | -44.18 | -12.93 | 6.23E-04 | *** |
| | 135 | PI-METHYLHISTIDINE | -30.06 | 8.03 | -45.92 | -14.20 | 7.69E-04 | *** |
| | 27 | TYROSINE_ACETAMINOPHEN_O-CRESOL_GLUTAMINE | 41.98 | 11.85 | 18.56 | 65.39 | 7.92E-04 | *** |
| | 8 | UNKNOWN | 14.83 | 4.22 | 6.49 | 23.16 | 8.79E-04 | *** |
| | 206 | GLUTAMATE | 71.22 | 20.52 | 30.68 | 111.76 | 0.001 | ** |
| | 13 | PHENYLALANINE_2-PHENYLPROPRIONATE | 21.29 | 6.15 | 9.13 | 33.44 | 0.001 | ** |
| | 207 | GLUTAMATE | 69.80 | 20.37 | 29.55 | 110.04 | 0.001 | ** |
| | 1 | FORMATE | 71.63 | 19.77 | 32.57 | 110.68 | 0.001 | ** |
| | 21 | TYROSINE | 27.23 | 8.05 | 11.33 | 43.14 | 0.001 | ** |
| | 203 | METHIONINE_GLUTAMINE_GLUTAMATE | -82.38 | 24.80 | -131.38 | -33.39 | 0.003 | ** |
| | 118 | UNKNOWN_GLYCEROL | 75.11 | 24.88 | 25.94 | 124.27 | 0.004 | ** |
| | 202 | GLUTAMINE_GLUTAMATE | -55.23 | 17.12 | -89.05 | -21.41 | 0.005 | ** |
| | 188 | PYRUVATE | 101.20 | 32.08 | 37.82 | 164.58 | 0.005 | ** |
| | 16 | PHENYLALANINE | 26.07 | 9.05 | 8.20 | 43.95 | 0.007 | ** |
| | 218 | LYSINE | -38.81 | 13.48 | -65.45 | -12.17 | 0.007 | ** |
| | 35 | UNKNOWN | 34.18 | 11.28 | 11.89 | 56.47 | 0.009 | ** |

| | | | | | | | |
|---|---|---|---|---|---|---|---|
| | 242 | ALLOISOLEUCINE_ISOLEUCINE | -53.71 | 19.91 | -93.06 | -14.37 | 0.012 | * |
| | 148 | UNKNOWN_BETAINE_GLUCOSE | 88.83 | 32.99 | 23.64 | 154.03 | 0.012 | * |
| | 246 | MOBILE LIPID -CH3 | -135.95 | 52.03 | -238.76 | -33.14 | 0.015 | * |
| | 152 | MOBILE LIPID -N(CH3)3 | -157.14 | 64.79 | -285.16 | -29.12 | 0.016 | * |
| | 34 | UNKNOWN | 22.33 | 9.14 | 4.26 | 40.40 | 0.024 | * |
| | 30 | UNKNOWN | 16.58 | 6.80 | 3.15 | 30.01 | 0.024 | * |
| | 174 | CITRATE | 41.36 | 17.34 | 7.09 | 75.62 | 0.028 | * |
| | 37 | MOBILE UNSATURATED LIPID | -60.15 | 27.57 | -114.63 | -5.67 | 0.031 | * |
| | 210 | GLYCYLPROLINE_ISOLEUCINE_UNKNOWN | -49.38 | 23.08 | -94.98 | -3.77 | 0.034 | * |
| | 129 | METHANOL | 49.14 | 23.34 | 3.03 | 95.25 | 0.037 | * |
| | 243 | ISOLEUCINE | -46.89 | 21.50 | -89.37 | -4.41 | 0.046 | * |
| | 145 | UNKNOWN | 20.74 | 23.04 | -24.79 | 66.27 | 0.370 | ns |
| Control vs Yes | 181 | GLUTAMINE | -147.28 | 30.02 | -206.60 | -87.96 | 3.60E-06 | **** |
| | 180 | GLUTAMINE | -97.34 | 19.90 | -136.66 | -58.02 | 3.82E-06 | **** |
| | 182 | GLUTAMINE | -150.45 | 31.05 | -211.80 | -89.10 | 4.68E-06 | **** |
| | 120 | GLYCEROL | 141.71 | 28.33 | 85.74 | 197.69 | 4.69E-06 | **** |
| | 147 | BETAINE | 408.25 | 83.75 | 242.77 | 573.73 | 8.24E-06 | **** |
| | 189 | GLUTAMATE_3-HYDROXYISOVALERATE | 134.21 | 27.55 | 79.78 | 188.65 | 8.36E-06 | **** |
| | 190 | GLUTAMATE | 133.93 | 27.64 | 79.31 | 188.56 | 9.38E-06 | **** |
| | 183 | GLUTAMINE | -97.51 | 20.85 | -138.70 | -56.32 | 9.63E-06 | **** |
| | 179 | GLUTAMINE | -52.37 | 11.57 | -75.22 | -29.52 | 1.81E-05 | **** |
| | 184 | GLUTAMINE | -58.42 | 13.04 | -84.18 | -32.67 | 2.19E-05 | **** |
| | 31 | UNKNOWN | 22.41 | 5.30 | 11.94 | 32.88 | 6.10E-05 | **** |
| | 3 | 4-PYRIDOXATE_ANSERINE | 19.62 | 4.62 | 10.50 | 28.74 | 1.12E-04 | *** |
| | 211 | ACETATE | 113.21 | 27.12 | 59.63 | 166.79 | 1.51E-04 | *** |
| | 227 | UNKNOWN | 146.56 | 37.59 | 72.29 | 220.83 | 2.18E-04 | *** |
| | 246 | MOBILE LIPID -CH3 | -214.82 | 52.76 | -319.07 | -110.58 | 2.26E-04 | *** |
| | 204 | GLUTAMINE_GLUTAMATE | -69.39 | 18.39 | -105.73 | -33.05 | 3.48E-04 | *** |
| | 185 | GLUTAMINE | -38.40 | 10.13 | -58.42 | -18.37 | 3.89E-04 | *** |
| | 13 | PHENYLALANINE_2-PHENYLPROPRIONATE | 24.43 | 6.24 | 12.10 | 36.75 | 4.08E-04 | *** |
| | 152 | MOBILE LIPID -N(CH3)3 | -256.53 | 65.70 | -386.34 | -126.71 | 4.27E-04 | *** |
| | 223 | LACTATE_THREONINE | 284.65 | 76.73 | 133.03 | 436.27 | 4.38E-04 | *** |
| | 199 | GLUTAMINE_2-METHYLGLUTARATE_AZELATE_SEBACATE | -31.74 | 8.59 | -48.71 | -14.77 | 4.61E-04 | *** |
| | 200 | GLUTAMINE_ACETAMINOPHEN | -55.23 | 14.98 | -84.84 | -25.63 | 4.77E-04 | *** |
| | 55 | LACTATE | 126.46 | 34.52 | 58.25 | 194.68 | 5.17E-04 | *** |
| | 134 | UNKNOWN | -30.01 | 8.02 | -45.86 | -14.17 | 6.23E-04 | *** |
| | 209 | UNKNOWN_GLUTAMATE | 85.68 | 23.92 | 38.41 | 132.96 | 6.91E-04 | *** |
| | 129 | METHANOL | 89.15 | 23.66 | 42.39 | 135.90 | 7.09E-04 | *** |
| | 8 | UNKNOWN | 16.11 | 4.28 | 7.65 | 24.56 | 7.19E-04 | *** |
| | 27 | TYROSINE_ACETAMINOPHEN_O-CRESOL_GLUTAMINE | 44.92 | 12.02 | 21.18 | 68.66 | 7.89E-04 | *** |

| | | | | | | | | |
|---|---|---|---|---|---|---|---|---|
| | 206 | GLUTAMATE | 72.26 | 20.80 | 31.16 | 113.37 | 0.001 | ** |
| | 210 | GLYCYLPROLINE_ISOLEUCINE_UNKNOWN | -85.72 | 23.41 | -131.97 | -39.48 | 0.001 | ** |
| | 218 | LYSINE | -49.86 | 13.67 | -76.88 | -22.84 | 0.001 | ** |
| | 207 | GLUTAMATE | 73.38 | 20.65 | 32.56 | 114.19 | 0.001 | ** |
| | 37 | MOBILE UNSATURATED LIPID | -100.21 | 27.96 | -155.46 | -44.97 | 0.001 | ** |
| | 21 | TYROSINE | 27.42 | 8.16 | 11.29 | 43.55 | 0.001 | ** |
| | 135 | PI-METHYLHISTIDINE | -26.81 | 8.14 | -42.89 | -10.73 | 0.002 | ** |
| | 1 | FORMATE | 65.78 | 20.04 | 26.17 | 105.38 | 0.002 | ** |
| | 148 | UNKNOWN_BETAINE_GLUCOSE | 112.59 | 33.46 | 46.48 | 178.70 | 0.003 | ** |
| | 242 | ALLOISOLEUCINE_ISOLEUCINE | -67.80 | 20.19 | -107.70 | -27.91 | 0.003 | ** |
| | 118 | UNKNOWN_GLYCEROL | 83.95 | 25.23 | 34.10 | 133.81 | 0.003 | ** |
| | 188 | PYRUVATE | 97.60 | 32.53 | 33.33 | 161.86 | 0.005 | ** |
| | 16 | PHENYLALANINE | 29.03 | 9.17 | 10.90 | 47.15 | 0.006 | ** |
| | 145 | UNKNOWN | 67.68 | 23.37 | 21.51 | 113.85 | 0.007 | ** |
| | 202 | GLUTAMINE_GLUTAMATE | -50.11 | 17.36 | -84.41 | -15.82 | 0.007 | ** |
| | 35 | UNKNOWN | 32.07 | 11.44 | 9.46 | 54.67 | 0.009 | ** |
| | 243 | ISOLEUCINE | -65.90 | 21.80 | -108.98 | -22.83 | 0.009 | ** |
| | 203 | METHIONINE_GLUTAMINE_GLUTAMATE | -65.86 | 25.14 | -115.54 | -16.18 | 0.015 | * |
| | 34 | UNKNOWN | 24.77 | 9.27 | 6.45 | 43.09 | 0.024 | * |
| | 30 | UNKNOWN | 18.44 | 6.89 | 4.83 | 32.06 | 0.024 | * |
| | 174 | CITRATE | 45.03 | 17.59 | 10.28 | 79.78 | 0.028 | * |
| Yes vs No | 145 | UNKNOWN | 46.94 | 14.34 | 18.61 | 75.27 | 0.004 | ** |
| | 129 | METHANOL | 40.01 | 14.52 | 11.32 | 68.70 | 0.010 | ** |
| | 246 | MOBILE LIPID -CH3 | -78.88 | 32.37 | -142.84 | -14.91 | 0.016 | * |
| | 152 | MOBILE LIPID -N(CH3)3 | -99.38 | 40.31 | -179.04 | -19.73 | 0.016 | * |
| | 210 | GLYCYLPROLINE_ISOLEUCINE_UNKNOWN | -36.35 | 14.36 | -64.72 | -7.97 | 0.019 | * |
| | 37 | MOBILE UNSATURATED LIPID | -40.06 | 17.16 | -73.96 | -6.16 | 0.031 | * |
| | 243 | ISOLEUCINE | -19.02 | 13.38 | -45.45 | 7.41 | 0.157 | ns |
| | 200 | GLUTAMINE_ACETAMINOPHEN | 12.20 | 9.19 | -5.97 | 30.36 | 0.187 | ns |
| | 218 | LYSINE | -11.05 | 8.39 | -27.63 | 5.52 | 0.190 | ns |
| | 190 | GLUTAMATE | 20.38 | 16.96 | -13.14 | 53.90 | 0.231 | ns |
| | 148 | UNKNOWN_BETAINE_GLUCOSE | 23.76 | 20.53 | -16.81 | 64.32 | 0.249 | ns |
| | 242 | ALLOISOLEUCINE_ISOLEUCINE | -14.09 | 12.39 | -38.57 | 10.39 | 0.257 | ns |
| | 203 | METHIONINE_GLUTAMINE_GLUTAMATE | 16.52 | 15.43 | -13.96 | 47.00 | 0.286 | ns |
| | 227 | UNKNOWN | -24.65 | 23.06 | -70.22 | 20.92 | 0.287 | ns |
| | 211 | ACETATE | 15.35 | 16.64 | -17.52 | 48.23 | 0.358 | ns |
| | 189 | GLUTAMATE_3-HYDROXYISOVALERATE | 15.52 | 16.90 | -17.88 | 48.92 | 0.360 | ns |
| | 223 | LACTATE_THREONINE | -39.38 | 47.08 | -132.41 | 53.65 | 0.404 | ns |
| | 147 | BETAINE | 42.23 | 51.39 | -59.30 | 143.77 | 0.412 | ns |
| | 13 | PHENYLALANINE_2-PHENYLPROPRIONATE | 3.14 | 3.83 | -4.42 | 10.70 | 0.413 | ns |

| | | | | | | | | |
|---|---|---|---|---|---|---|---|---|
| | 3 | 4-PYRIDOXATE_ANSERINE | 2.05 | 2.83 | -3.55 | 7.65 | 0.470 | ns |
| | 179 | GLUTAMINE | 4.78 | 7.10 | -9.24 | 18.81 | 0.501 | ns |
| | 135 | PI-METHYLHISTIDINE | 3.25 | 4.99 | -6.62 | 13.11 | 0.517 | ns |
| | 199 | GLUTAMINE_2-METHYLGLUTARATE_AZELATE_SEBACATE | 3.42 | 5.27 | -7.00 | 13.83 | 0.518 | ns |
| | 31 | UNKNOWN | -2.04 | 3.25 | -8.46 | 4.39 | 0.532 | ns |
| | 120 | GLYCEROL | 10.05 | 17.38 | -24.29 | 44.39 | 0.564 | ns |
| | 118 | UNKNOWN_GLYCEROL | 8.84 | 15.48 | -21.75 | 39.43 | 0.569 | ns |
| | 55 | LACTATE | -11.48 | 21.18 | -53.33 | 30.38 | 0.589 | ns |
| | 209 | UNKNOWN_GLUTAMATE | -7.83 | 14.68 | -36.84 | 21.17 | 0.594 | ns |
| | 16 | PHENYLALANINE | 2.95 | 5.63 | -8.17 | 14.08 | 0.600 | ns |
| | 8 | UNKNOWN | 1.28 | 2.63 | -3.91 | 6.47 | 0.626 | ns |
| | 202 | GLUTAMINE_GLUTAMATE | 5.12 | 10.65 | -15.92 | 26.16 | 0.631 | ns |
| | 1 | FORMATE | -5.85 | 12.30 | -30.15 | 18.45 | 0.635 | ns |
| | 30 | UNKNOWN | 1.87 | 4.23 | -6.49 | 10.22 | 0.660 | ns |
| | 204 | GLUTAMINE_GLUTAMATE | 4.84 | 11.28 | -17.45 | 27.14 | 0.668 | ns |
| | 34 | UNKNOWN | 2.44 | 5.69 | -8.80 | 13.68 | 0.669 | ns |
| | 27 | TYROSINE_ACETAMINOPHEN_O-CRESOL_GLUTAMINE | 2.94 | 7.37 | -11.63 | 17.51 | 0.691 | ns |
| | 174 | CITRATE | 3.67 | 10.79 | -17.65 | 24.99 | 0.734 | ns |
| | 35 | UNKNOWN | -2.12 | 7.02 | -15.99 | 11.75 | 0.763 | ns |
| | 134 | UNKNOWN | -1.46 | 4.92 | -11.18 | 8.26 | 0.767 | ns |
| | 207 | GLUTAMATE | 3.58 | 12.67 | -21.46 | 28.62 | 0.778 | ns |
| | 180 | GLUTAMINE | 2.55 | 12.21 | -21.58 | 26.67 | 0.835 | ns |
| | 188 | PYRUVATE | -3.60 | 19.96 | -43.03 | 35.83 | 0.857 | ns |
| | 185 | GLUTAMINE | -1.00 | 6.22 | -13.28 | 11.29 | 0.873 | ns |
| | 206 | GLUTAMATE | 1.04 | 12.77 | -24.18 | 26.27 | 0.935 | ns |
| | 184 | GLUTAMINE | -0.59 | 8.00 | -16.40 | 15.21 | 0.941 | ns |
| | 181 | GLUTAMINE | 1.26 | 18.42 | -35.14 | 37.66 | 0.946 | ns |
| | 21 | TYROSINE | 0.18 | 5.01 | -9.72 | 10.08 | 0.971 | ns |
| | 183 | GLUTAMINE | -0.41 | 12.79 | -25.68 | 24.86 | 0.974 | ns |
| | 182 | GLUTAMINE | -0.28 | 19.05 | -37.93 | 37.36 | 0.988 | ns |
